# BRIDGE prioritizes human and mouse gene correspondences using functional and disease evidence

**DOI:** 10.64898/2026.05.10.720506

**Authors:** Jinyun Yan, Ze Cao

## Abstract

Selecting mouse genes to model human disease requires evidence that a perturbation represents the biological function of interest. Orthology provides an evolutionary reference, but does not by itself resolve disease-related functional differences between species. We present BRIDGE, a framework that combines species-labelled disease associations and functional annotations through graph representations, optimal transport and rank fusion to prioritize human–mouse gene correspondences. The rankings include non-orthologous candidates and distinguish alternative effectors, distributed signalling modules and multiple human genes linked to one mouse counterpart. BRIDGE achieved 61.8–66.7% Recall@5 across three functional-correspondence benchmarks. In 25 curated drug-translation cases, 15 parsed mouse targets ranked first; a separate phenotype-informed screen retained 54 of 533 candidates in the final processed dataset. These results organize gene selection around the function a disease experiment must interrogate and nominate hypotheses for individual or combined perturbations. Because the benchmarks partly reuse annotation evidence, retrieval does not establish phenotypic or pharmacological equivalence. BRIDGE provides a basis for reviewing mouse genes before disease-model construction or drug screening, and for investigating how disease-associated functions are distributed across mammalian genes. These applications require disease-specific experimental validation.

## Introduction

Which mouse gene should be perturbed to investigate a human disease mechanism? Orthology provides the evolutionary starting point by identifying genes related through speciation [1,2]. Yet loss of corresponding human and mouse genes can have different phenotypic consequences [3]. A disease experiment therefore requires an additional judgement: whether the chosen mouse perturbation represents the function relevant to the question. Depending on that function, an informative counterpart could be the ortholog, a paralog, another functional analogue or a group of interacting genes.

The intended experiment determines what counts as a useful match. Studying a disease-associated pathway requires the relevant process to operate in the chosen cells and tissues. Testing a therapeutic also requires the appropriate molecular interaction. Similar sequences alone establish neither conserved expression nor ligand recognition or downstream signalling.

Conversely, a mouse gene may provide an informative pathway readout without binding a drug directed against the human target. Gene selection and pharmacological model selection thus overlap, but they are not the same decision.

Comparative methods already connect species at several biological levels. Orthology resources describe ancestry, and single-cell methods such as CAME and SATURN align cellular identities and molecular representations [1,2,4,5]. PhenoDigm and subsequent cross-species phenotype analyses already connect animal-model phenotypes with human diseases [6,7]. These capabilities leave a more specific model-design question: which mouse gene or gene set should be considered when a one-to-one ortholog inadequately represents a human gene’s disease-associated role? We use non-orthologous to distinguish a candidate from the relevant orthology assignment, not to imply that it lacks shared ancestry. Many alternatives remain members of the same ancestral gene family.

Established biology suggests that this question requires more than a single type of mapping. Mouse chemokine ligands and their shared receptor distribute neutrophil recruitment across a signalling module [8]. Pentraxin acute-phase responses differ between species [9,10], whereas human CASP4 and CASP5 connect to a related mouse inflammatory-caspase system with distinct stimulus-recognition properties [11,12]. Such relationships change both the perturbations to compare and the meaning of a negative result. Regulatory subfunctionalization and mutation-induced compensation further motivate testing how duplicated genes divide or retain a process [13,14].

To organize this choice, we developed BRIDGE to rank human–mouse correspondences from disease-associated and functional evidence (Figure 1). The framework connects species-labelled disease and phenotype associations to Gene Ontology, pathway and interaction information, then combines graph representations, optimal transport and rank fusion. We first ask what these rankings recover under different definitions of correspondence. Retrospective cases then distinguish the biological interpretations of a match, and phenotype evidence narrows the candidates for model review. The output is a set of genes and gene sets to test, including non-orthologous alternatives, rather than a declaration of biological equivalence.

**Figure 1.**
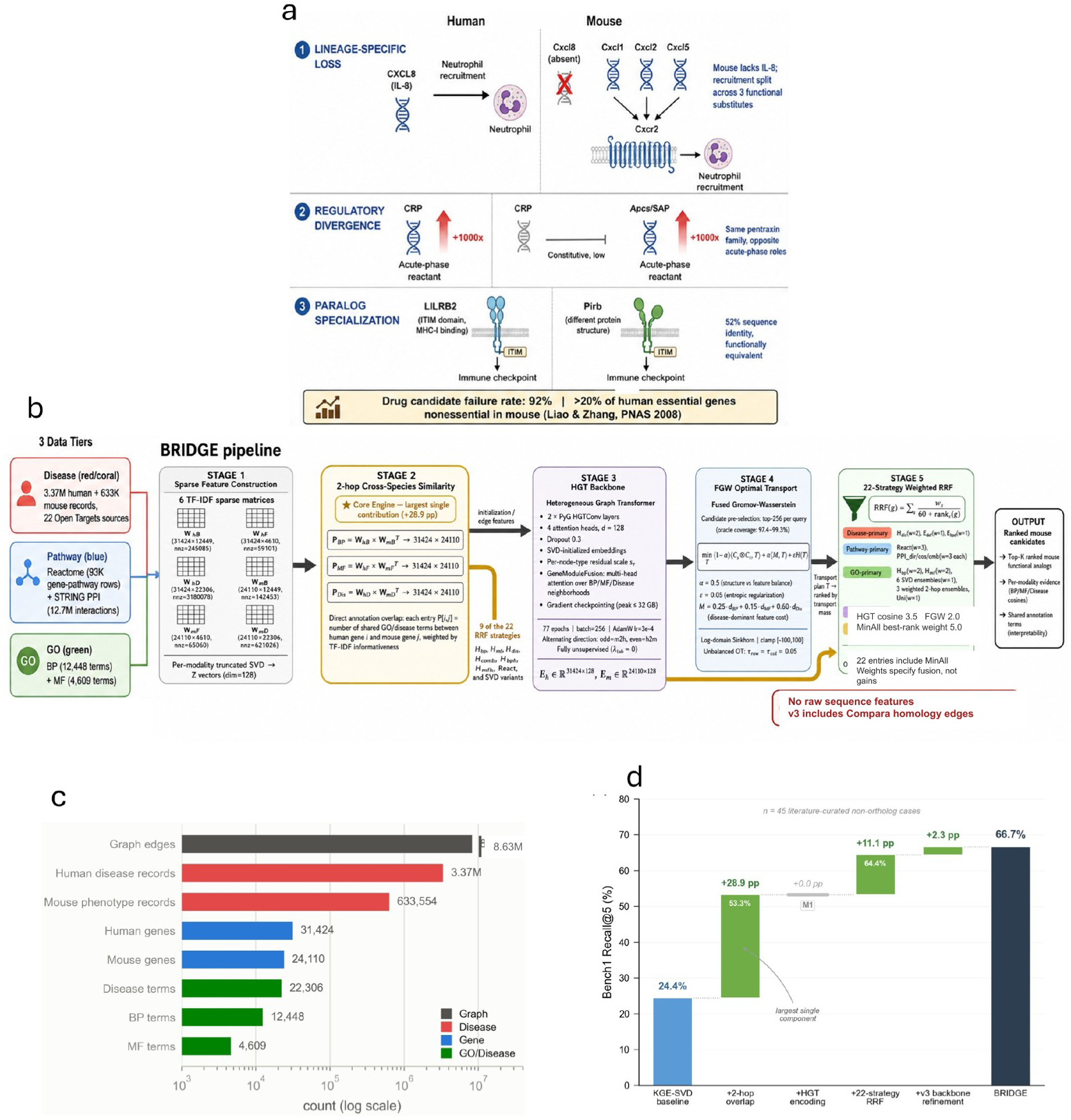
Disease-model questions and the BRIDGE workflow. (a) Literature-based examples distinguish ancestry from disease-relevant function. They are not new experimental validations, and the schematic drug-attrition percentage is not an outcome measured here. (b) Species-labelled associations feed graph representation, FGW and reciprocal rank fusion. The final diagram specifies FGW weight 2.0, HGT-cosine weight 3.5 and 22 fusion entries including MinAll. V3 includes Compara homology information although raw sequence features are not used. (c) The graph inventory contains 94,897 nodes and 8,625,606 forward/reverse annotation edges before additional v3 relations. (d) The Bench1 progression is a configuration-level comparison. Its two-hop value is 53.3%, distinct from the standalone value in Figure 2b. Graph inventory and the training trace are shown in Figure 1—figure supplement 1 and Figure 1—figure supplement 2, respectively.

## Results

### Functional retrieval depends on the correspondence being tested

BRIDGE retrieved functional correspondences in benchmarks whose answers were not primarily defined by one-to-one orthology. Recall@5 was 66.7% on literature-curated Bench1 and 61.8% on Bench2-HARD (Figure 2a,b; Tables 1 and 2), and 64.8% on annotation-derived FuncBench (Supplementary Files 2, 3 and 7). Ensembl Compara recovered almost all Bench2-EASY assignments but none in Bench2-HARD. Because the hard subset specifically selected poorly Ensembl-ranked pairs, that contrast tests a different correspondence objective; it is not evidence that orthology inference generally fails.

**Figure 2.**
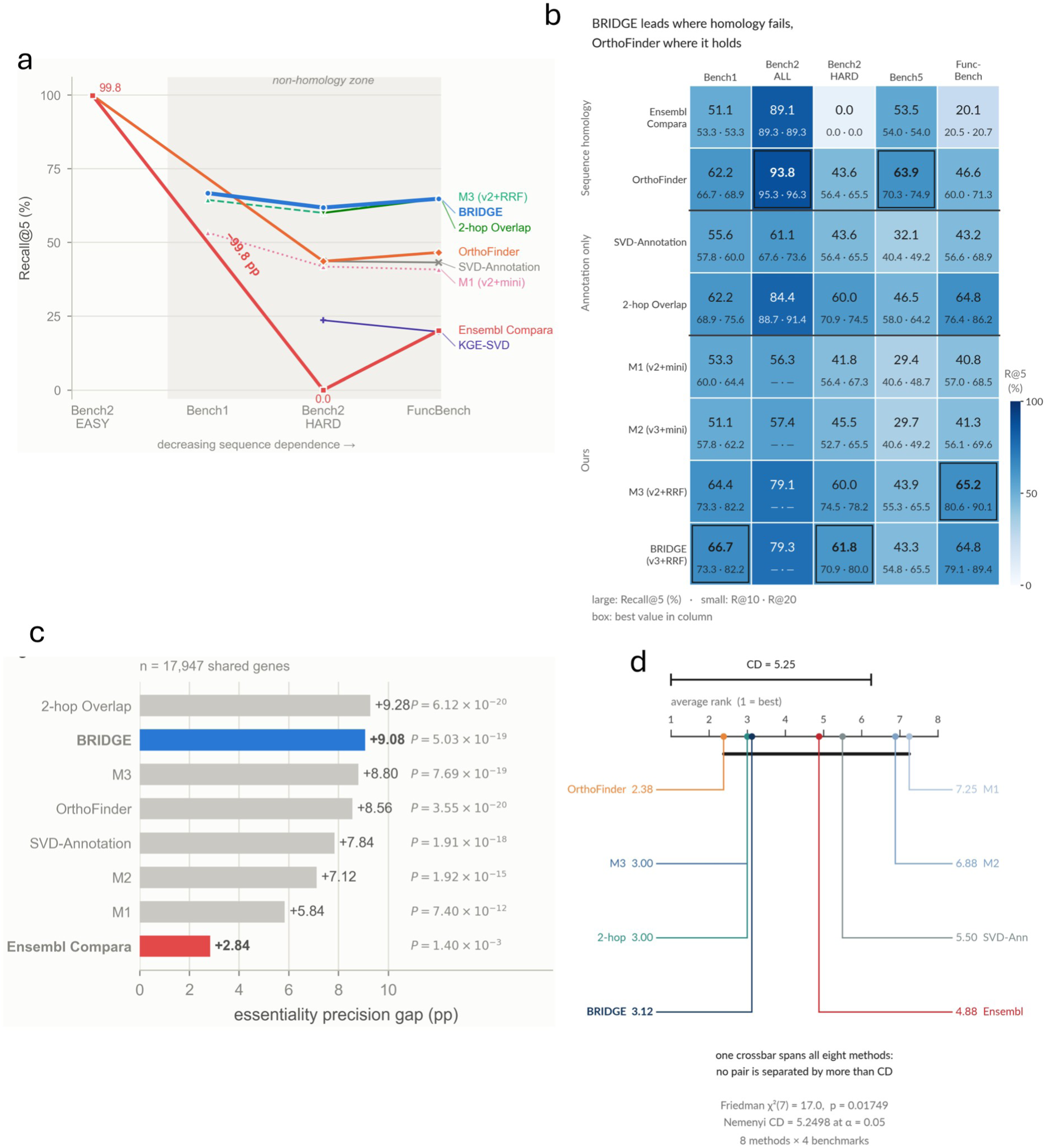
Retrieval depends on the definition of correspondence. (a) Recall@5 across benchmarks with different dependence on orthology-based labels. M4 achieves 61.8–66.7% on Bench1, Bench2-HARD and FuncBench. Bench2-HARD explicitly selects poorly Ensembl-ranked pairs. (b) Recall@5 with Recall@10 and Recall@20 annotations; dashes denote unavailable combinations. Bench2-ALL uses 512 pairs (Supplementary Files 2 and 7). (c) Essentiality-concordance gap in a shared-gene universe of 17,947 genes. The sampled query comparison is distinguished from that universe in Supplementary File 2. M4 has a 9.08-percentage-point gap, with reported one-sided Mann–Whitney P = 5.03 × 10⁻¹⁹; the Ensembl gap is 2.84 points, P = 1.40 × 10⁻³. These are within-method group comparisons, not a superiority test between methods. (d) Friedman χ²(7) = 16.985, P = 0.0175; Nemenyi critical difference 5.25 across eight methods and four benchmarks. No method pair exceeds that difference. Standalone component results and the complete performance matrix are shown in Figure 2—figure supplement 1 and Figure 2—figure supplement 2, respectively.

**Table 1.**
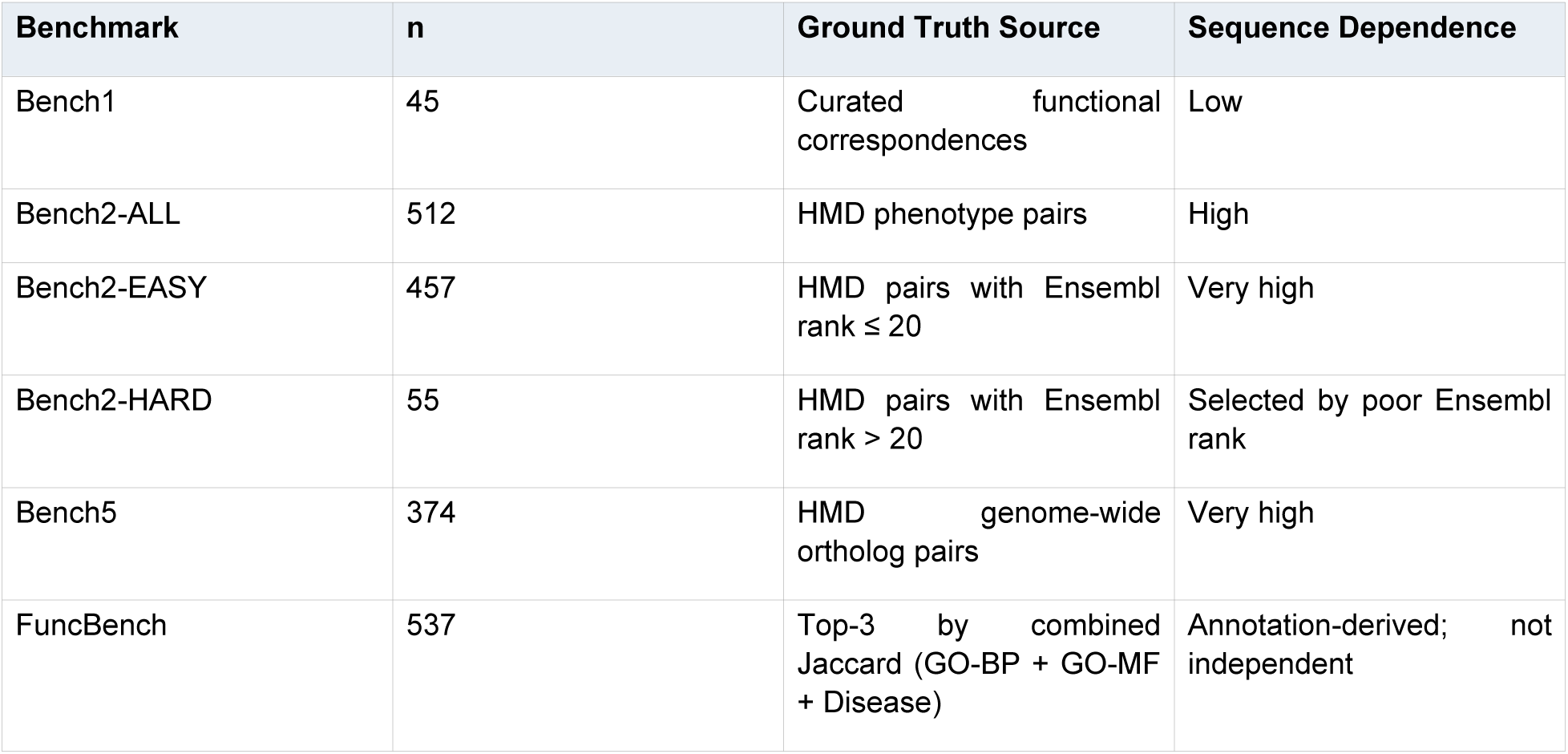
Benchmark-design overview. Six benchmarks distinguish functional retrieval from orthology recovery. Bench2-HARD selects poorly Ensembl-ranked pairs; FuncBench defines answers by combined annotation Jaccard. The sequence-dependence categories describe benchmark construction, not absence of homology information from the model inputs.

| Benchmark | n | Ground Truth Source | Sequence Dependence |
| --- | --- | --- | --- |
| Bench1 | 45 | Curated functional correspondences | Low |
| Bench2-ALL | 512 | HMD phenotype pairs | High |
| Bench2-EASY | 457 | HMD pairs with Ensembl rank $\leq 20$ | Very high |
| Bench2-HARD | 55 | HMD pairs with Ensembl rank $> 20$ | Selected by poor Ensembl rank |
| Bench5 | 374 | HMD genome-wide ortholog pairs | Very high |
| FuncBench | 537 | Top-3 by combined Jaccard (GO-BP + GO-MF + Disease) | Annotation-derived; not independent |

**Table 2.** Cross-benchmark Recall@5 summary (%). Best entry per column in bold.

| Method | Bench1 | Bench2-ALL | Bench2-HARD | Bench5 | FuncBench | Bench3 Gap |
| --- | --- | --- | --- | --- | --- | --- |
| Ensembl Compara | 51.1 | 89.1 | 0.0 | 53.5 | 20.1 | +2.84 |
| Ensembl-SVD | 62.2 | **93.8** | 43.6 | **63.9** | 46.6 | +8.56 |
| SVD-Annotation | 55.6 | 61.1 | 43.6 | 32.1 | 43.2 | +7.84 |
| 2-hop Overlap | 62.2 | 84.4 | 60.0 | 46.5 | 64.8 | **+9.28** |
| M1 (v2+mini) | 53.3 | 56.3 | 41.8 | 29.4 | 40.8 | +5.84 |
| M2 (v3+mini) | 51.1 | 57.4 | 45.5 | 29.7 | 41.3 | +7.12 |
| M3 (v2+RRF) | 64.4 | 79.1 | 60.0 | 43.9 | **65.2** | +8.80 |
| BRIDGE | **66.7** | 79.3 | **61.8** | 43.3 | 64.8 | +9.08 |

No retrieval strategy was strongest under every definition of correspondence. Orthology-oriented methods remained strongest on orthology-rich benchmarks, while BRIDGE and direct annotation-overlap methods were competitive on functional benchmarks. Across eight methods and four benchmarks, no pair exceeded the Nemenyi critical difference (Figure 2d).

An essentiality comparison provided a separate measure of concordance: BRIDGE showed a 9.08-percentage-point precision gap between essential and nonessential query groups (Figure 2c; Supplementary Files 2 and 7). This association does not establish that any retrieved gene reproduces a particular human disease phenotype.

The resulting capability is annotation-informed prioritization within the tested settings. Some benchmark targets draw on the same kinds of functional annotations used as inputs, and v3 includes homology information (Figure 1b). The fused Gromov–Wasserstein (FGW) component is additionally restricted by candidate pre-selection, whereas the full ensemble includes mouse-genome-wide ranking lists. Separate benchmark definitions and component results make these dependencies explicit (Supplementary Files 3 and 6).

### Retrieved correspondences suggest different perturbation designs

A distributed correspondence points to a module-level experiment rather than a single replacement gene. For CXCL8, the illustrated ranking contained Cxcl5, Cxcl1, Cxcl2 and Cxcr2 at positions 1–4 (Figure 3a). These ligands and their shared receptor offer distinct perturbations of neutrophil recruitment, consistent with their context-dependent roles in mouse studies [8]. They are not interchangeable molecules. The illustrated configuration also differs from the separately scored Ortholog-Override Criterion (OOC) evaluation, so the two sets of ranks are kept separate (Supplementary File 4).

**Figure 3.**
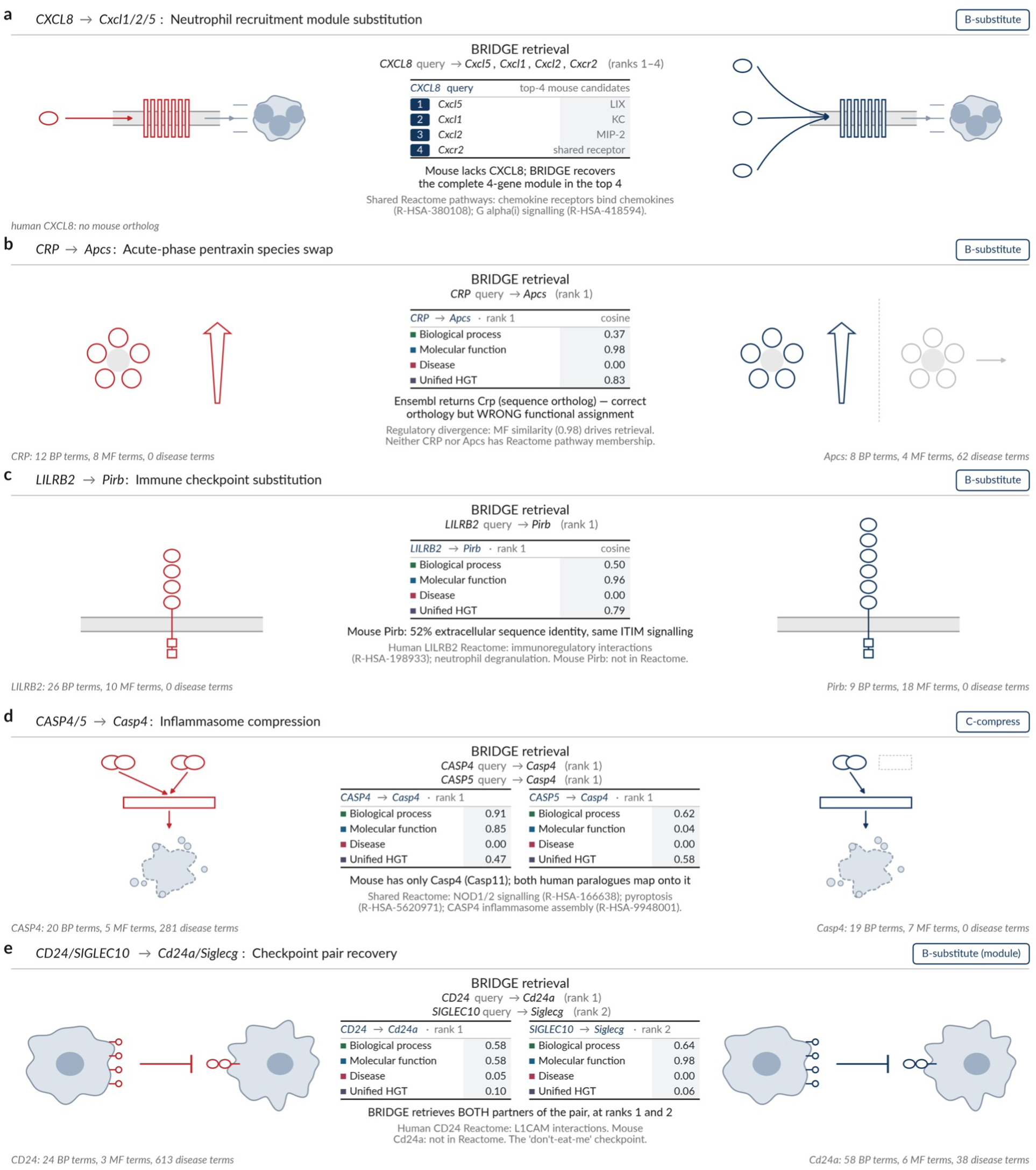
Alternative effectors and conserved modules. (a) The illustrated CXCL8 candidates are Cxcl5, Cxcl1, Cxcl2 and Cxcr2 at ranks 1–4 [8]. (b) CRP–Apcs and (c) LILRB2–Pirb are displayed at rank 1 [9,10,15]. (d) Human CASP4 and CASP5 each retrieve mouse Casp4 at rank 1 [11,12]. (e) CD24–Cd24a and SIGLEC10–Siglecg are displayed at ranks 1 and 2 [16,17]. These retrospective examples illustrate distinct correspondence patterns. Figure-specific ranks and OOC-run ranks are identified separately in Supplementary File 4. The CXCL8 illustration and OOC query use different configurations; the SIGLEC10 label in the module illustration is also kept separate from its individual-query OOC rank. OOC results and anchor-case phenotype comparisons are provided in Figure 3—figure supplement 1 and Figure 3—figure supplement 2, respectively.

Individual matches instead nominated particular mouse readouts. CRP retrieved Apcs at rank 1 with molecular-function cosine similarity of 0.98 (Figure 3b), consistent with serum amyloid P component’s contribution to the mouse acute-phase response [9,10]. LILRB2 retrieved Pirb at rank 1 (Figure 3c), recovering an established relationship in inhibitory signalling and amyloid-associated biology [15]. These examples support reviewing the relevant function of each candidate, not assuming that the conventional counterpart is inactive or that all biochemical activities are conserved.

Many-to-one and paired correspondences raised different design questions. Both human CASP4 and CASP5 mapped to mouse Casp4 in the inflammatory-caspase example (Figure 3d). The shared pathway relationship coexists with differences in stimulus recognition [11,12]. In the CD24–SIGLEC10 example, Cd24a and Siglecg occupied ranks 1 and 2, respectively, within a paired regulatory axis (Figure 3e) [16,17]. The curated relationships and case-level evaluations are retained separately in Supplementary Files 3 and 4.

Together, the cases distinguish alternative effectors, distributed modules and multiple human genes converging on one mouse counterpart. These patterns organize hypotheses about what to perturb and how to interpret the response. They remain retrospective recoveries of established relationships. None of four scored cases passed the stricter A-promote evaluation (Figure 3—figure supplement 1), so the results do not establish that BRIDGE generally replaces a suitable ortholog with a superior paralog.

### Drug-related retrieval complements orthology without establishing pharmacological equivalence

The drug-related cases chiefly supported consideration of conventional counterparts. Among 25 curated drug-translation cases, 15 parsed mouse targets ranked first, and 14 of those were one-to-one orthologs (Figure 4a; Supplementary File 5). Functional ranking therefore complemented orthology instead of consistently selecting alternatives. Unscorable targets and missing comparisons remain in the complete roster. The endpoint was target retrieval, not drug efficacy or successful clinical translation.

**Figure 4.**
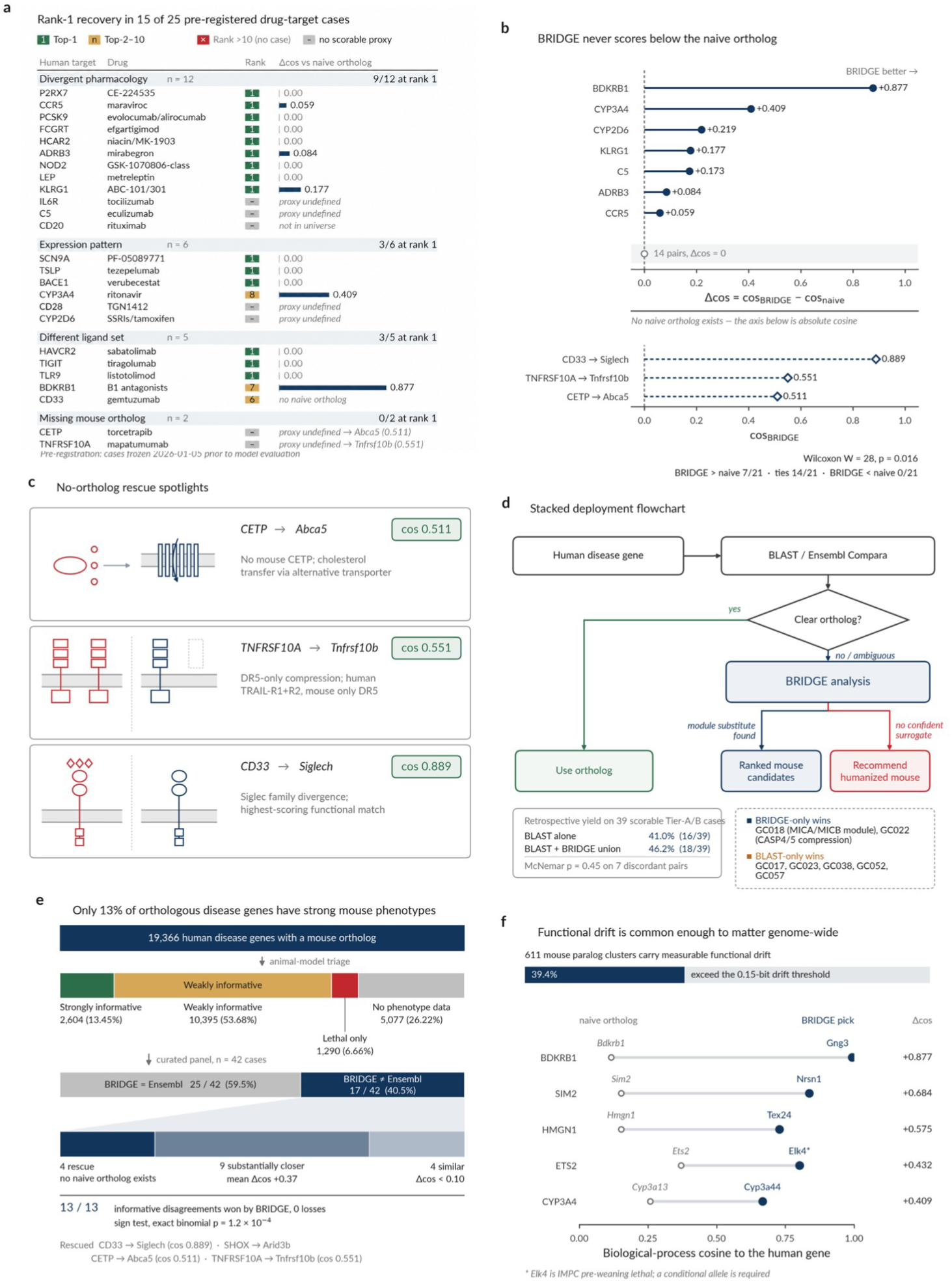
Drug-related retrieval and functional interpretation. (a) The 25 cases comprise 12 divergent-pharmacology, six expression-pattern, five ligand-set and two missing-ortholog classifications; 15 parsed mouse targets rank first (Supplementary File 5). (b) Twenty-one paired BP comparisons contain seven positive and fourteen zero differences. Selection maximizes the same score, so the displayed Wilcoxon annotation is not independent evidence of M4 superiority. (c) CETP–Abca5, TNFRSF10A–Tnfrsf10b and CD33–Siglech are BP-nearest associations. (d) The 39-case OOC union gives baseline, M4 and union pass rates of 41.0%, 33.3% and 46.2%. The baseline is labelled BLAST in the plot; its study implementation is described in Methods. (e) Resource triage and the 42-case BP comparison have different denominators; the latter displays 25 unchanged and 17 alternative assignments. (f) Among 611 paralog clusters, 241 exceed a mean Jensen–Shannon divergence of 0.15 bit. Annotation divergence does not establish evolutionary changes in function. The last-digit difference in the ETS2 comparison between panels 4f and 6d reflects display rounding. The union analysis, annotation-divergence analysis and CD33 comparison are shown in Figure 4—figure supplements 1–3, respectively.

An annotation-level comparison nominated additional candidates without independently testing their quality as models. Seven of 21 paired cases had a positive biological-process (BP) cosine difference from the Ensembl comparator; the remaining fourteen were tied (Figure 4b; Supplementary File 5). The candidate was selected by maximizing the same BP score used to calculate that difference. These descriptive comparisons identify pathway relationships for review, but cannot establish better pharmacological models.

CD33 shows why the identity of the ranked candidate matters. Siglech was the BP-nearest candidate, whereas the drug-case M4 rank of 6 belonged to the parsed target Cd33, a different target and candidate context (Figure 4c; Figure 4—figure supplement 3; Supplementary File 5). Siglec-H signals through DAP12 rather than intrinsic ITIMs [18]. Shared annotations consequently establish neither CD33-like signalling nor recognition by an anti-CD33 therapeutic. The comparison proposes an immune-regulatory experiment, not a validated replacement for a human-target pharmacology assay.

Combining mappings produced only modest complementarity in the separate 39-case OOC analysis. The displayed union pass rate was 46.2%, versus 41.0% for the baseline and 33.3% for M4 (Figure 4d; Figure 4—figure supplement 1). The exact McNemar test did not establish a difference (P = 0.45). These results support reviewing both mappings while leaving improvement in model selection unestablished. The case-level OOC endpoint is distinct from ordinary rank-1 retrieval (Supplementary File 4).

### Annotation overlap and ranking composition account for substantial retrieval signal

The component analysis placed the retrieval results in context. In the Bench1 ablation, adding two-hop annotation overlap increased Recall@5 from 24.4% to 53.3% (Figure 5a). Subsequent graph encoding did not improve every benchmark, whereas the full ensemble improved the v2 model over its mini-ensemble counterpart across the three displayed functional benchmarks. Direct annotation structure and ensemble composition therefore supplied substantial signal; graph learning and optimal transport alone do not explain the results.

**Figure 5.**
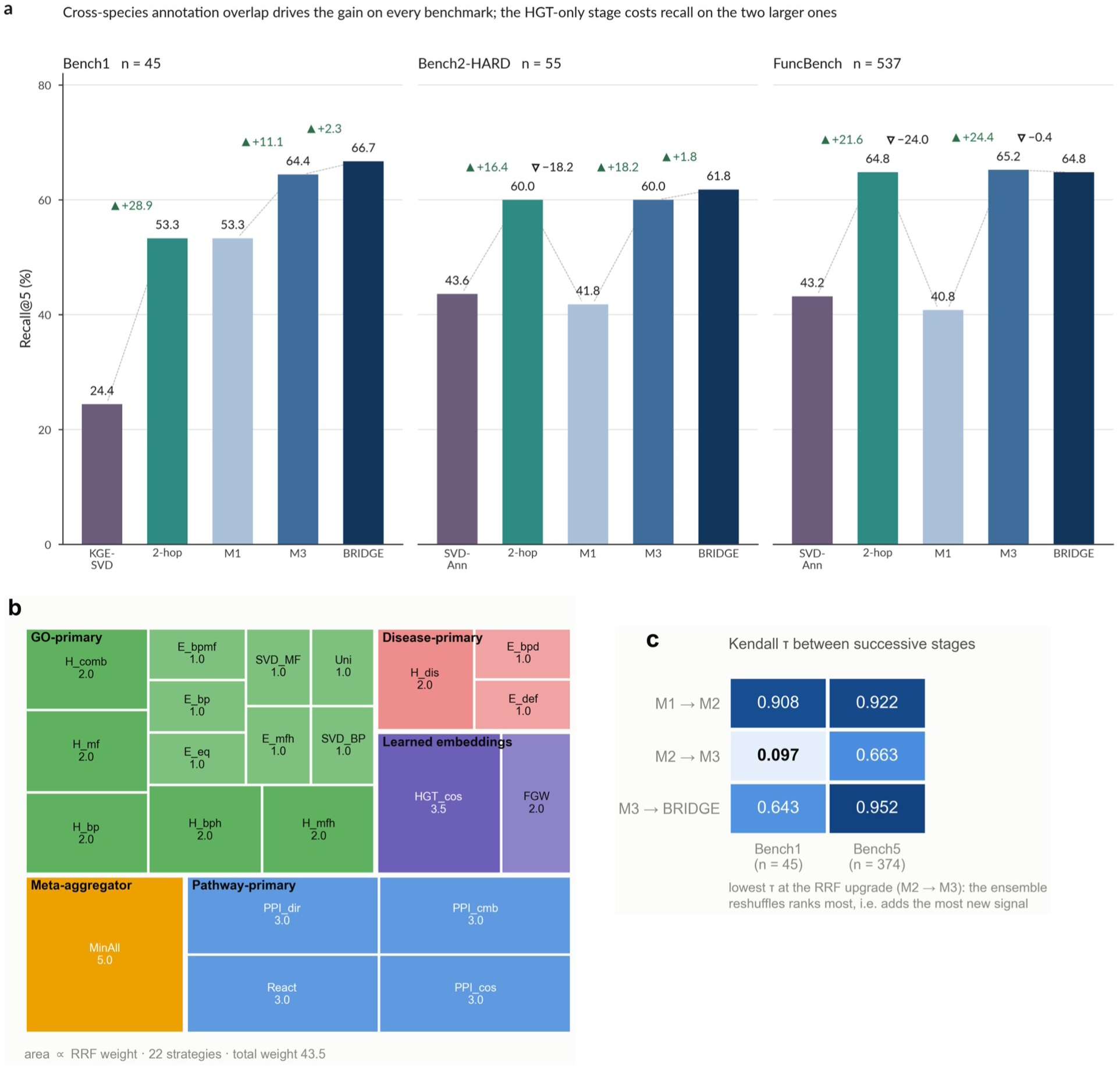
Component comparisons and fusion configuration. (a) Recall@5 on Bench1 (n = 45), Bench2-HARD (n = 55) and FuncBench (n = 537). The baseline, two-hop, M1, M3 and M4 sequences are 24.4, 53.3, 53.3, 64.4 and 66.7%; 43.6, 60.0, 41.8, 60.0 and 61.8%; and 43.2, 64.8, 40.8, 65.2 and 64.8%, respectively. The Bench1 two-hop ablation value differs from the standalone 62.2% in Figure 2b. (b) The displayed fusion includes FGW weight 2.0 and HGT-cosine weight 3.5. Weight magnitude does not measure an isolated accuracy contribution. (c) Kendall-τ comparisons for M1/M2, M2/M3 and M3/M4 on Bench1 and Bench5. The largest reordering coincides with a change in both backbone and retrieval configuration. Supplementary Files 1, 2, 6 and 7 preserve the distinct analyses. FGW parameter sensitivity is shown in Figure 5—figure supplement 1.

The final ensemble combined disease-, pathway- and GO-derived rankings with learned similarities (Figure 5b). FGW and HGT-cosine weights were 2.0 and 3.5, respectively, specifying the contribution of ranking lists rather than causal effects on biology or accuracy. The standalone performance matrix and ablation used different two-hop configurations, with Bench1 values of 62.2% and 53.3%, respectively. Their per-case strategy, component and cross-benchmark analyses remain distinct (Supplementary Files 1, 6 and 7).

### Phenotype evidence identifies a smaller set for disease-specific review

A functional match is informative for model selection only in the context of available phenotype evidence. Resource triage classified 2,604 of 19,366 genes as strong-informative under the stated criteria (Figure 6a). The remaining categories were weak-informative, lethal-only and without phenotype data. This classification describes recorded evidence, not a genome-wide experimental failure rate: a missing phenotype record leaves the relevant function unresolved.

**Figure 6.**
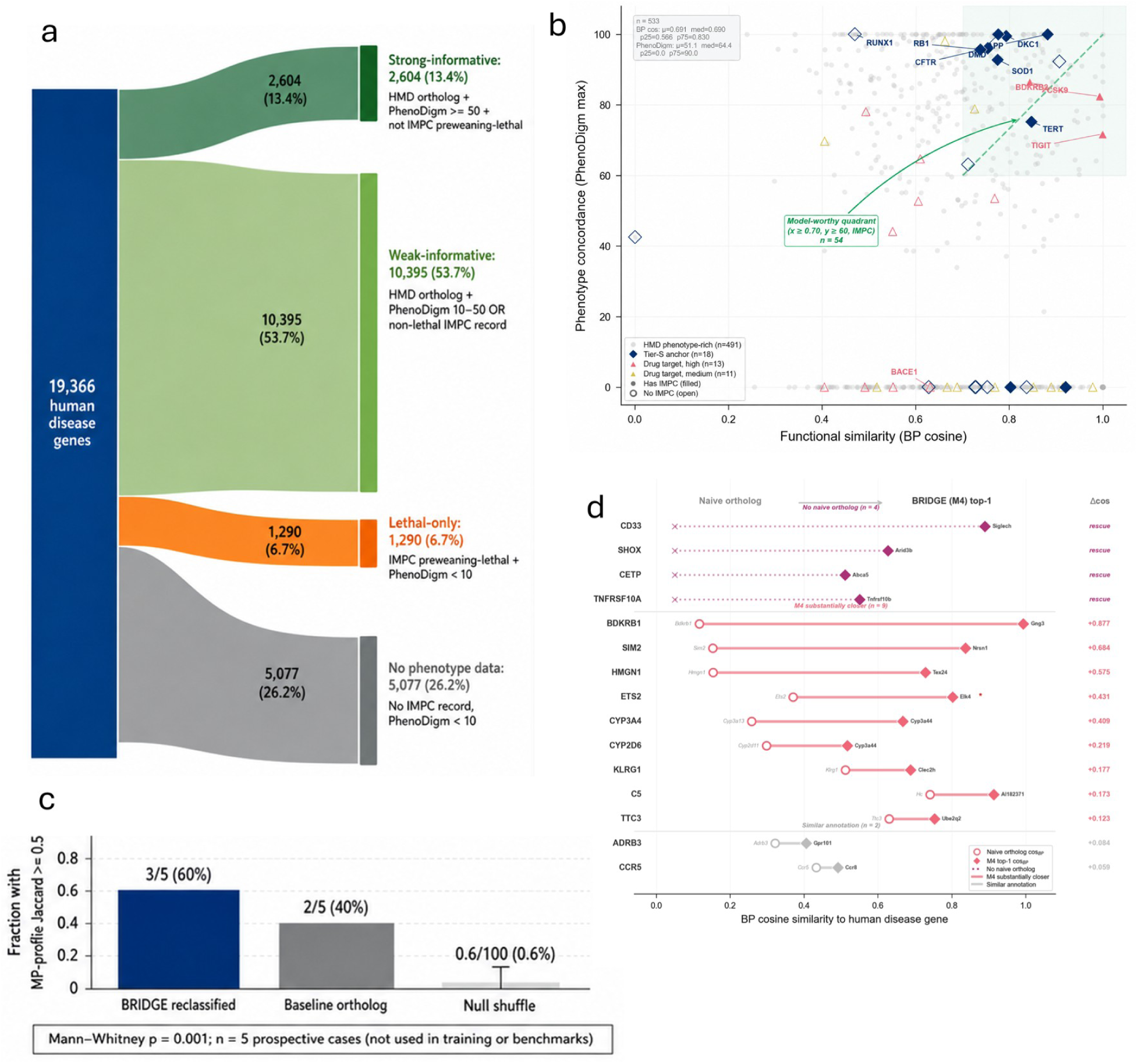
Phenotype-informed candidate prioritization. (a) Of 19,366 genes, 2,604 are strong-informative, 10,395 weak-informative, 1,290 lethal-only and 5,077 without phenotype data. (b) The final processed plotting dataset contains 533 candidates; 54 meet BP cosine ≥0.70, maximum PhenoDigm score ≥60 and IMPC-record presence (Supplementary File 8). This is a candidate screen, not experimental validation. (c) The five-case exploratory panel reports three BRIDGE and two comparator assignments meeting its displayed phenotype-profile Jaccard criterion (≥0.5); its aggregate does not establish general model quality. (d) The illustrated candidate comparisons are descriptive. This panel uses a different case set from Figure 4e. CD33–Siglech and the drug-case Cd33 rank concern different target/candidate contexts (Supplementary File 5). Four additional candidate correspondences are illustrated in Figure 6—figure supplement 1.

Applying functional and phenotype criteria together reduced the candidate set to 54 of 533 entries in the final processed plotting dataset (Figure 6b; Supplementary File 8). All three criteria were required: BP cosine of at least 0.70, maximum PhenoDigm score of at least 60 and an IMPC record. This is a prioritization screen, not a validation endpoint. In particular, the maximum PhenoDigm score may describe a disease-model match other than the human disease of interest, making disease-specific review necessary.

The shortlist and annotation records support that next review step (Supplementary Files 8 and 9). Four illustrations nominate CFH–Cfhr4, C1S–C1s2, CYP2B6–Cyp2b19 and REG1A– Reg3d, with BP, MF and disease evidence shown separately (Figure 6—figure supplement 1). The broader catalogues are not a one-to-one inventory of those illustrations. Likewise, the five-case phenotype-profile panel and reclassification comparison supply exploratory context (Figure 6c,d), not independent evidence that a nominated model predicts human treatment response.

## Discussion

The practical contribution of BRIDGE is to make the choice of mouse perturbation explicit. A gene can be a close evolutionary counterpart without answering every disease-model question, and a functionally related candidate can be informative without being an equivalent molecule. Taken together, the retrieval and case analyses support comparing these possibilities within one evidence framework. They do not support routinely discarding orthology: the appropriate correspondence depends on the process and intervention being studied.

### Separate pathway modelling from target pharmacology

For preclinical research, the immediate use is candidate review before animal-model selection. Disease associations and functional annotations can highlight inflammatory, metabolic or immune-regulatory readouts when gene loss, regulatory divergence or family specialization complicates the conventional mapping. Such a comparison helps formulate what the mouse experiment is intended to measure. The curated drug-related cases illustrate this reasoning process; they provide no test of clinical-efficacy prediction.

A useful next step is to ask which part of the proposed intervention the candidate can represent. Similar downstream processes may justify a pathway experiment even if the mouse protein lacks the expression pattern or binding site needed for a human-specific therapeutic. Conversely, the ortholog may remain the relevant choice for target-engagement studies despite another gene’s closer annotation match. Tissue expression, ligand recognition, molecular interactions and pharmacology therefore need their own evaluation. The CD33 example makes this separation concrete.

### Use correspondence patterns to formulate combinatorial experiments

A distributed match changes the unit of perturbation under consideration. If several mouse genes contribute to the process associated with one human gene, loss of a single member may leave substantial activity intact. The CXCL8-related module consequently motivates comparisons among individual ligands, selected combinations and receptor perturbation. These designs test different mechanisms rather than alternative implementations of a ranking. Each needs a defined disease-related phenotype and a hypothesis about why the selected components should matter.

A many-to-one match poses the opposite problem: a single mouse perturbation may combine functions separated between human paralogs. Perturbing Casp4 could affect processes associated with both CASP4 and CASP5, limiting attribution to either human gene. More generally, a candidate set can contain redundant effectors, sequential pathway components or genes sharing only broad annotations. Allele- and mutant-transcript-dependent genetic compensation adds another source of variation [13]. Viability, tissue specificity and developmental timing remain essential design constraints.

These possibilities are experimental questions, not outputs already predicted by BRIDGE. The framework neither optimizes gene combinations nor infers epistasis or combined-knockout phenotypes, and this study generated no new multi-gene knockout animals. Its proposed contribution is to narrow the candidates for single and combined perturbation. Comparisons against orthology- and pathway-guided designs, evaluated with phenotypes withheld during development, would test whether that prioritization yields more informative models.

### Distinguish evolutionary hypotheses from annotation divergence

The correspondence patterns also suggest how to ask about the organization of function across mammals. A process may remain associated with an ortholog, become partitioned among paralogs or involve another interacting module. The human–mouse cases and 611-cluster annotation-divergence analysis nominate families in which to examine these possibilities (Figure 4f; Figure 4—figure supplement 2). However, uneven annotation coverage can resemble functional divergence. Neither a rank nor an annotation difference establishes the direction or mechanism of evolutionary change.

Resolving that ambiguity requires alignment scores to be considered alongside gene trees, duplication histories and tissue-resolved functional evidence. Regulatory subfunctionalization of Hox1 genes illustrates why coding-sequence relationships alone may not explain the distribution of present-day functions [14]. Broader mammalian comparisons could then test whether a disease-associated process is conserved while the contributing genes change. This remains prospective because the present evaluation covers only human and mouse.

### Test candidate value with evidence independent of ranking inputs

Dependence between inputs and evaluation labels limits what the benchmarks establish. Disease and GO annotations are uneven and incomplete, and related annotations define some benchmark targets. Orthology-derived mappings and homology information in v3 add further dependence, precluding a sequence-free or wholly orthology-independent interpretation. In addition, the figure-labelled published methods are study-specific proxies built from this study’s embeddings or transport outputs, not independent software executions (Supplementary Files 6 and 7). The comparisons therefore characterize this retrieval framework rather than establish superiority over reproduced external methods.

The next evaluation must test whether candidate priorities change the biological information obtained. Selected retrospective cases, a small prospective phenotype-profile panel and thresholding of curated processed data do not answer that question under blinded conditions. Candidate pre-selection and manually specified fusion weights also affect the rankings. A discriminating test would compare functional, orthology-based and combined selection using experimental phenotypes unavailable during development. Claims about pharmacological model choice would additionally require independent drug-response experiments.

Disease-informed functional alignment can organize the evidence behind a model-design decision without resolving that decision automatically. BRIDGE makes alternative genes and gene sets available for comparison with conventional orthologs, linking candidate review to explicit perturbation hypotheses. Its proposed uses in combinatorial models and mammalian evolution follow from those patterns, but remain to be tested. The decisive criterion is whether disease-specific experiments show that a nominated correspondence improves the design or interpretation of the mouse model.

## Methods

BRIDGE ranks cross-species candidates by combining a heterogeneous graph with graph-based representation, fused Gromov–Wasserstein transport and weighted reciprocal rank fusion [19,20,21]. GO and disease terms are shared between species. The final configuration has 22 fusion entries, including MinAll (Figures 1b and 5b). The per-case strategy comparison in Supplementary File 1 is a separate analysis, not an evaluation of the final M4 configuration.

### Data sources and preprocessing

#### Functional annotations

GO biological-process (BP) and molecular-function (MF) associations for human and mouse were obtained through Gene Ontology and Ensembl [1,22]. After processing, the representation contained 245,085 human BP, 59,101 human MF, 142,453 mouse BP and 65,060 mouse MF associations (Figure 1—figure supplement 1). These are nonzero entries in the processed representation, not the earlier raw annotation counts. The displayed term inventory excluded root terms.

#### Disease associations

We assembled human gene–disease associations from Open Targets and represented mouse associations through phenotype-related mappings [23,24]. Separate species-labelled edges identify the origin of each association. These labels do not remove dependence introduced by orthology-derived mappings, either between species or between input annotations and evaluation labels.

Bench2-HARD comprises HMD pairs for which the Ensembl comparator rank exceeds 20. FuncBench defines the top three partners using the mean of BP, MF and disease binary Jaccard scores, subject to its annotation-similarity threshold. The supplied case roster does not document systematic exclusion of every orthology-transferred label. Accordingly, FuncBench evaluates internal annotation retrieval rather than independent experimental validation (Supplementary File 3).

#### Species-separated evidence architecture

Separate edge types connect human and mouse genes to shared GO and disease nodes. The processed representation has 245,085 human BP, 142,453 mouse BP, 59,101 human MF, 65,060 mouse MF, 3,180,078 human disease and 621,026 mouse disease nonzero entries. The corresponding count-derived BP means are 7.80 per human gene and 5.91 per mouse gene, with human BP matrix density approximately 0.00063 (Figure 1—figure supplement 1). These graph summaries are not independent counts of biological observations.

#### Auxiliary signals

Pathway-based rankings use Reactome memberships, and interaction-based rankings use STRING associations [25,26].

#### Sparse feature construction

For each (gene, term) pair, we populated six sparse matrices with the raw GO count or, for disease associations, the Open Targets overall association score:

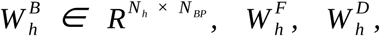

and the three mouse counterparts 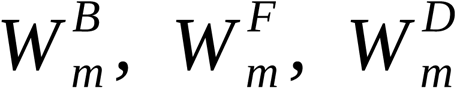.

We winsorised disease scores at quantile q = 0.995, rescaled them linearly to [0, 1], and applied TF-IDF column weighting followed by row ℓ2 normalisation. GO matrices underwent the same TF-IDF and row-ℓ2 processing but no winsorisation.

#### Final graph statistics

The final figure inventory contains 31,424 human genes, 24,110 mouse genes, 12,448 BP terms, 4,609 MF terms and 22,306 disease terms, for a total of 94,897 nodes. Across the six annotation matrices, 4,312,803 nonzero entries generate 8,625,606 forward and reverse annotation edges (Figure 1c; Figure 1—figure supplement 1). Additional cross-species relations specified for v3 are not included in that annotation-edge total.

### Unified heterogeneous graph construction

We constructed one heterogeneous graph in which both species share GO and disease nodes, rather than constructing and subsequently aligning separate species graphs. Let

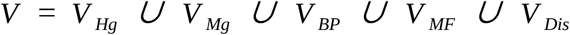

denote the five node types. The production backbone v2 has 12 edge types: six forward bipartite relations (HG–BP, HG–MF, HG–Dis, MG–BP, MG–MF, MG–Dis) and their six reverse counterparts. A GO or disease node t can connect genes from both species, allowing message passing to exchange cross-species features without direct gene-to-gene edges.

For v3, explicit cross-species relations supplement the shared-term graph. The Compara homology information shown in Figure 1b is part of v3; omitting raw sequence features therefore does not make the graph homology-free. Annotation-derived pairwise similarities provide further retrieval evidence:

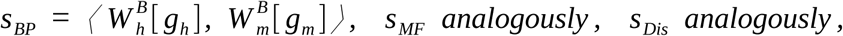

The modality-specific similarities feed cross-species ranking channels. These channels are distinct from the graph relations, and annotation overlap alone does not establish orthology. Comparing v2/v3 thus changes the available cross-species information rather than isolating a purely sequence-independent intervention.

### HGT backbone

The HGT backbone [27] uses L = 2 PyG HGTConv layers [28], H = 4 attention heads, hidden dimension d = 128 and dropout 0.3. We initialise each node type’s learnable embedding from a modality-specific truncated SVD of its sparse feature matrix, with truncation ranks

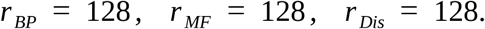

For each human or mouse gene, concatenating three 128-dimensional modality vectors produces a 384-dimensional vector before projection:

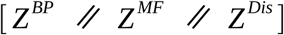

We project the concatenated gene vectors and the term-node features to the same hidden dimension of 128. An edge-weight blending term (κ = 0.1) incorporates the original TF-IDF weight into the attention logits. Each layer produces an additive residual,

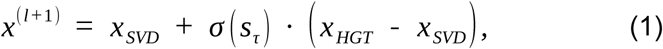

with a learnable scale sτ for each node type, initialised to

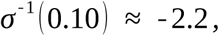

which keeps the start of training close to the SVD solution. Each HGTConv call uses gradient checkpointing, with peak memory below 32 GB.

To construct the candidate-selector prior Π0, we rank all mouse genes for each human gene gh using the blended SVD retrieval score

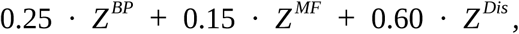

and retain the top-256 targets. We then apply MMR diversification [29] (

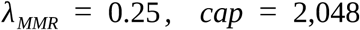

) before renormalising the scores into a probability vector. No supervised benchmark labels are used to construct this prior. The final workflow uses batches of 256 source genes (Figure 1b), with each batch’s candidate target set combining its pre-selected lists. Non-gold targets within that set participate through the restricted coupling, not a separate explicit negative-sampling step.

GeneModuleFusion forms per-modality queries over each gene’s BP, MF and Disease neighbourhoods and combines them through multi-head attention. Its entropy regulariser has coefficient 0.001. Human and mouse have separate fusion modules, each using mean-centring for isotropy.

#### Training objective

For a mini-batch of B source genes and their associated candidate target sets, we optimise the composite loss

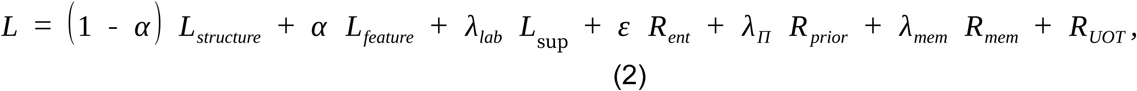

where ℒstructure and ℒfeature are the FGW structure and feature costs defined below; Rent denotes Sinkhorn entropy; Rprior penalises KL divergence from the candidate-selector prior Π0; Rmem anchors the coupling to an exponential-moving-average memory bank of previous couplings; and RUOT groups the unbalanced-OT marginal KL penalties. The supervised term remains disabled throughout, with λlab = 0 (fully unsupervised setting). The coefficients are α = 0.5, ε = 0.05, λΠ = 0.5 and λmem = 0.3.

#### Optimisation

The final workflow uses AdamW [30] with learning rate 3 × 10⁻⁴, batch size 256 and 77 training epochs (Figure 1b). The displayed trace has its best checkpoint at epoch 73 (Figure 1—figure supplement 2); the 40-epoch marker is a reference point, not the training endpoint. This trace does not document a shared optimization history for all ablation variants.

#### Training protocol, hardware, and reproducibility

The reported software environment is Python 3.11, PyTorch 2.3.1+cu121 [31] and PyTorch Geometric 2.7.0 [28]. The archived v2 hardware record names an NVIDIA GeForce RTX 4070 Ti SUPER. The evaluations compare methods and benchmarks, not repeated random-seed runs, and no multi-seed variance is reported. Reproduction therefore requires the evaluation-specific configuration, checkpoint and processed-data version.

### Fused Gromov–Wasserstein optimal-transport alignment

We use the trained gene embeddings Eh ∈ ℝN_h × d, Em ∈ ℝN_m × d, together with per-modality SVD features

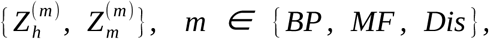

to retrieve the top-256 mouse genes for each human query through the candidate selector’s blended score

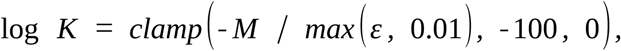

The FGW candidate selector applies MMR diversification [29] with lambda = 0.25 and cap 2,048. Estimated candidate coverage is 97.8% for Bench1, 99.1% for Bench2-ALL, 99.3% for Bench2-EASY, 97.4% for Bench2-HARD, 99.0% for Bench5 and 98.2% for FuncBench. These estimates concern transport pre-selection. Because full RRF also incorporates mouse-genome-wide rankings, its Recall@k is not universally conditional on membership in this restricted FGW pool.

We solve the fused Gromov–Wasserstein problem [19,21] over the restricted |I| × |J| coupling,

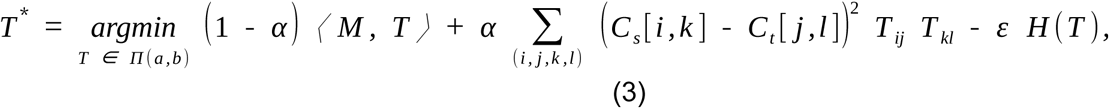

where M weights the BP, MF and disease feature costs by 0.25, 0.15 and 0.60, respectively. These are specified hyperparameters rather than estimates of each modality’s biological importance. Cs and Ct contain cosine-distance structure costs, and (a, b) define the transported mass. We set α = 0.5 and entropic regularisation ε = 0.05, use row and column KL penalties τrow = τcol = 0.05, and warm-start from the candidate-selector prior.

#### Numerically stabilised Sinkhorn

For numerical stability, we implement Sinkhorn in the log domain [21] with

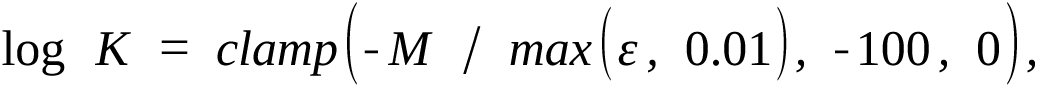

#### and apply the usual dual updates

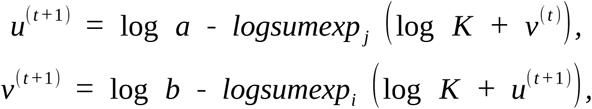

while clamping each potential to [−100, 100] and limiting optimization to 100 iterations. Non-finite couplings are replaced by the independence coupling a ⊗ b. In the FGW parameter panel, Bench5 Recall@5 is 7.42% at α = 0.5 and ε = 0.05; the grid maximum is 7.69% across 364 scorable pairs out of 374 (Figure 5—figure supplement 1). The selected parameter cell is therefore not the grid optimum.

### Weighted reciprocal rank fusion

Weighted reciprocal rank fusion [20] combines the active rankings shown in Figures 1b and 5b. For mouse candidate g, rank_s(g) denotes its one-based rank under strategy s, and w_s the assigned strategy weight. We calculate the fused score as

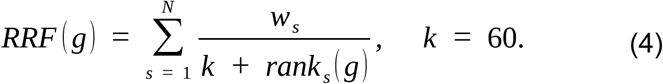

We sort candidates by decreasing RRF score using rank constant k = 60. The final weights are 2.0 for FGW, 3.5 for HGT cosine, 5.0 for MinAll, 3.0 for Reactome and 3.0 for each PPI list. Weights were manually specified, without an independent optimization study. The separate 53-case strategy analysis in Supplementary File 1 reports full22 ranks, which are distinct from the final M4 ranks.

#### Tier 1; disease-primary

The disease-primary tier includes H_dis (two-hop on the disease bipartite graph; w = 2.0), E_def (SVD ensemble with wDis = 0.60) and E_bpd (balanced BP + Disease ensemble). The FGW feature cost also emphasizes disease through wDis = 0.60.

#### Tier 2; pathway-primary

The pathway-primary tier includes React (Reactome two-hop; w = 3.0) and three STRING-PPI variants: PPI_dir, PPI_cos and PPI_cmb, each with w = 3.0.

#### Tier 3; GO-primary

The GO and learned channels comprise BP and MF two-hop similarities, SVD rankings, combined annotation rankings, HGT embedding cosine and FGW transport mass. MinAll takes each candidate’s best rank across the contributing strategies. The displayed fusion configuration, legacy strategy ablation and candidate annotation scores in Supplementary File 9 describe different quantities and are not substituted for one another.

### Model variants for ablation

The ablation variants pair backbone and ranking configurations as follows: M1 uses v2 with the mini ensemble, M2 uses v3 with the mini ensemble, M3 uses v2 with full RRF, and M4 (BRIDGE) uses v3 with full RRF. These contrasts alter the backbone, retrieval-layer composition or both. Supplementary Files 2, 6 and 7 report method-level metrics, standalone component results and cross-benchmark summaries. A contrast changing several inputs does not isolate the contribution of one component.

### Ortholog-Override Criterion

OOC applies subtype-specific rules. A-promote passes when a curated non-ortholog target ranks strictly above the naive one-to-one ortholog; a case lacking the required comparison remains unscored. B-substitute requires a gold rank of 1 and, when a distinct naive ortholog exists, an ortholog rank exceeding 10. C-compress requires each human input to retrieve a gold member at rank 1. Finding a finite gold rank alone therefore does not establish a pass when no ortholog is found.

The OOC roster includes 57 curated cases and 25 drug cases. In the separately scored M4 evaluation, gold ranks are available for 71 cases and pass/fail outcomes for 46. The displayed subtype outcomes are 0/4 for A-promote, 11/36 for B-substitute and 2/6 for C-compress (Figure 3—figure supplement 1). Supplementary File 4 distinguishes these OOC-run ranks from the different ranking configuration illustrated in Figure 3; the configurations are not pooled in the aggregate. The union analysis uses its own 39-case subset. Invalid archived subtype entries for CCR5, SCN9A and KLRG1 leave their pass/fail outcomes unreported. These records remain unscored rather than being reclassified from curated hypotheses.

### Benchmarks and comparison methods

The figure evaluation comprises Bench1 (45 cases), Bench2-ALL (512 pairs), Bench2-EASY (457 pairs), Bench2-HARD (55 pairs), Bench5 (374 pairs) and FuncBench (537 pairs). The Bench2-ALL values in Supplementary Files 2 and 7 refer to the final 512-pair cohort. Unavailable metrics remain missing rather than being imported from the earlier 509-pair evaluation. All candidates come from a universe of 24,110 mouse genes.

The pair-level evidence roster for Bench1, Bench2-HARD, FuncBench and OOC is provided in Supplementary File 3. Bench1 has 45 pair-level cases, and FuncBench has 537 pairs from 184 evaluable human queries. The essentiality figure’s 17,947 shared genes define a universe distinct from the archived sample of 500 essential and 500 nonessential human queries. Supplementary File 2 retains these denominators separately.

We compared Ensembl Compara, SVD-Annotation, 2-hop Overlap and the figure-labelled comparators. In this study, OrthoFinder and BLAST label Ensembl-prioritized SVD retrieval; CAME labels v3 HGT-cosine retrieval; SATURN labels FGW-only retrieval; and GenePlexusZoo labels unified-SVD cosine retrieval. These are internal proxies, not executions of the published software [4,5,32]. Additional v2/v3 embedding, FGW-only and unified-SVD component comparisons are reported in Supplementary File 6.

### Statistical tests

The reported essentiality analysis uses a one-sided Mann–Whitney comparison between essential and nonessential query groups. Friedman and Nemenyi summaries compare eight methods across four benchmarks, with no pair exceeding the critical difference. BP-nearest-minus-comparator differences are descriptive because selection maximizes the score being compared. Drug-case Wilcoxon and reclassification sign tests are not independent evidence of model superiority. The phenotype screen reports a threshold count rather than a significance test: in the final processed dataset, candidates must have BP cosine ≥0.70, maximum PhenoDigm score ≥60 and an IMPC record (Supplementary File 8) [6,33]. No repeated-seed uncertainty estimate is available.

### Supporting data and reproducibility

Supplementary Files 1–9 supply the strategy analysis, benchmark results and case evidence, OOC evaluation, drug-case roster, component comparisons, performance summary, prospective and phenotype-screening records, and candidate annotation evidence. Reproduction additionally requires the processed resource versions, the configuration-to-checkpoint mapping and the analysis scripts.

## Supporting information

Supplementary File 1 - Ranking strategy ablation

Supplementary File 2 - Benchmark and essentiality results

Supplementary File 3 - Benchmark case evidence

Supplementary File 4 - OOC case results

Supplementary File 5 - Drug translation cases

Supplementary File 6 - Internal component baselines

Supplementary File 7 - Cross-benchmark summary

Supplementary File 8 - Prospective shortlist and phenotype screen

Supplementary File 9 - Candidate annotation evidence

## Use of generative artificial intelligence

Claude 4.6 (Anthropic) and Codex (OpenAI) assisted with manuscript preparation, language revision, table organization and consistency checks. This revision harmonized author-designated processed data and figure versions; it did not rerun model training or generate new biological measurements. The authors are responsible for verifying data provenance, computational results, scientific interpretation and references.

## Data availability

Processed tabular results accompanying this manuscript are provided in Supplementary Files 1–9. The study draws on public annotation and phenotype resources cited in Methods.

## Code availability

The BRIDGE source code is publicly available at https://github.com/AmaranthZe/BRIDGE.

## Author contributions

Ze Cao: Conceptualization, Methodology, Software, Supervision. Jinyun Yan: Formal analysis, Visualization, Writing – original draft.

## Supplementary files

Supplementary File 1. Per-case ranking strategies and fusion ablations. Fifty-three cases with individual strategy ranks and full22, core5 and single-best summaries. This historical strategy analysis is distinct from the final M4 benchmark and OOC evaluations. Values of 24,110 in restricted outputs are sentinels.

Supplementary File 2. Benchmark and essentiality results. Twenty-four method-by-benchmark records. Bench2 uses the final 512-pair figure cohort; unsupported metrics from the earlier 509-pair cohort are not carried forward. Retrieval recalls are percentages, MRR is dimensionless, essentiality precision and hit-rate fields retain the source scale, and essentiality gaps are percentage points. The sampled 500-essential plus 500-nonessential query comparison is distinguished from the 17,947-gene shared universe. These are not random-seed repetitions.

Supplementary File 3. Benchmark case evidence. The 878 pair-level records comprise 45 Bench1, 55 Bench2-HARD, 537 FuncBench and 241 expanded OOC records. Human and mouse identifiers, benchmark memberships, curated relationship types and available annotation evidence are retained. An empty literature or orthology-transfer-removal field means unavailable information, not a negative finding or verified exclusion.

Supplementary File 4. Case-level OOC evaluation. The 820 records comprise 82 cases evaluated under ten stored method labels. Human inputs and curated mouse target sets are linked by case identifier. OOC_run_best_gold_rank is the stored evaluation rank; illustrated_best_gold_rank identifies the different rank shown in Figure 3 where applicable. The illustration and individual-query OOC contexts must not be pooled as one run. The module-hit field is labelled K20, matching the implemented threshold. Published-method labels identify the archived study comparisons described in Methods, not independently reproduced external packages. Invalid archived subtype entries for CCR5, SCN9A and KLRG1 are left blank; their pass/fail fields remain unreported, and Supplementary File 5 subtype hypotheses are not substituted for an executed evaluation.

Supplementary File 5. Drug-translation cases and evaluated candidates. Twenty-five curated cases with literature context, parsed ranking target, M4 target rank, BP-nearest candidate and Ensembl comparison. A curated suspected proxy, a parsed gold target and a BP-nearest candidate are different roles. Missing comparisons are retained. The TGN1412 and niacin/MK-1903 rows correspond to CD28 and HCAR2, respectively, matching Figure 4a. Candidate retrieval does not establish drug response or pharmacological interchangeability.

Supplementary File 6. Internal component baselines. Twenty records spanning Bench1, Bench2-ALL, Bench2-EASY, Bench2-HARD and FuncBench for HGT_standard_v2, HGT_crossspec_v3, FGW_OT_only and KGE_SVD. These are component-level comparisons, not independent reproductions of externally named published methods. Recall fields are percentages and MRR is dimensionless. Sample sizes and missing values are retained from the corresponding evaluation. The separate Bench5 values in Figure 2— figure supplement 1 are not tabulated here.

Supplementary File 7. Cross-benchmark performance summary. Sixteen rows combine the eight-method Recall@5 summary and the eight-method functional/sequence-cliff summary. Bench2-ALL Recall@5 uses the final 512-pair figure values. B3_gap is measured in percentage points and MRR is dimensionless. This table contains no IEA sensitivity experiment. The standalone Bench1 two-hop value follows Figure 2b and is not silently substituted for the distinct ablation-stage value in Figures 1d and 5a.

Supplementary File 8. Prospective shortlist and phenotype-screening records. The record_type field separates 30 prospective_shortlist records from 533 phenotype_screen records. The shortlist score and the screen’s BP cosine are different quantities. Screening rows contain the final processed plotting dataset. Fifty-four rows satisfy BP cosine ≥0.70, maximum PhenoDigm score ≥60 and IMPC-record presence. The score summarizes the maximum available phenotype match and is not disease-specific validation. Blank fields are not zeros. This table contains 563 records in total.

Supplementary File 9. Candidate-level annotation evidence. The 316 human–mouse candidate records include BP, MF and disease cosine similarities, overlap measures and shortlist membership. weighted_annotation_score is the supplied 0.25 BP + 0.15 MF + 0.60 disease score, not an RRF score. catalogue_order retains the supplied candidate ordering, which need not descend by that score. illustrated_rank preserves supplied case-specific ranks, including records for POT1 and APOL1 that are no longer shown in the current four-case supplement. illustrated_panel links only records displayed in the current figure; C1S candidates refer to Figure 6—figure supplement 1b. Blank panel references do not mean that the underlying candidate record or rank is missing. No unreported ranks are inferred.

