## Supplementary figures and legends for "BRIDGE prioritizes human and mouse gene correspondences using functional and disease evidence"


Jinyun Yan[1] and Ze Cao[2]

[1] Department of Bioengineering, Sichuan University, Chengdu 610065, China
[2] Department of Molecular Biology, Princeton University, Princeton, NJ 08544, USA


### Figure 1—figure supplement 1

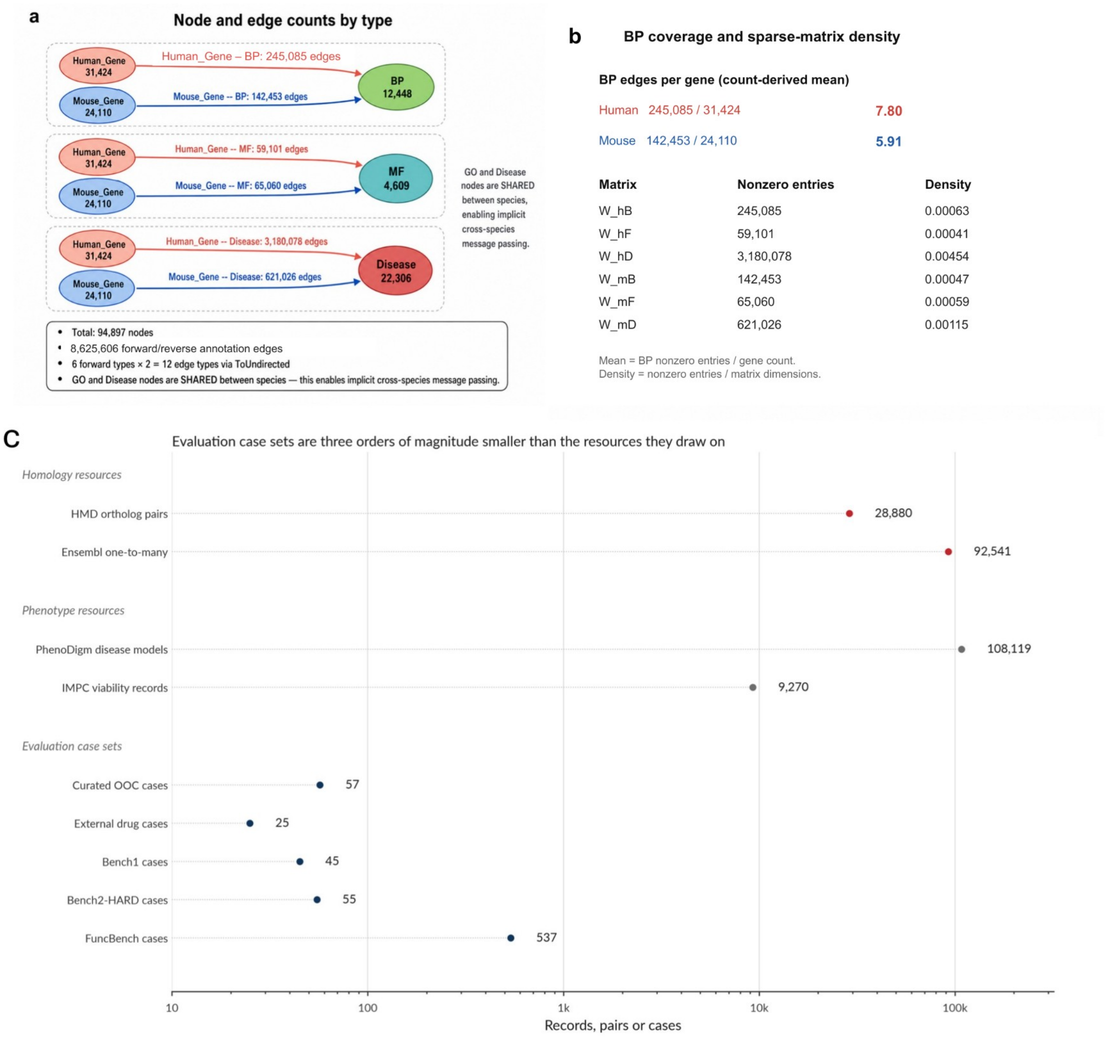


Figure 1—figure supplement 1. Graph inventory and annotation coverage. (a) The final graph inventory contains 94,897 nodes, including 31,424 human genes, 24,110 mouse genes, 12,448 BP terms, 4,609 MF terms and 22,306 disease terms. Human and mouse BP associations number 245,085 and 142,453. The six annotation relations and their reverses contain 8,625,606 edges. (b) Count-derived BP means are 7.80 per human gene and 5.91 per mouse gene; the human BP sparse-matrix density is approximately 0.00063. These are count summaries, not a kernel-density estimate. (c) Resource inventories and evaluation-set sizes describe different units. The curated OOC and drug rosters contain 57 and 25 cases, respectively.

Figure 1—figure supplement 2

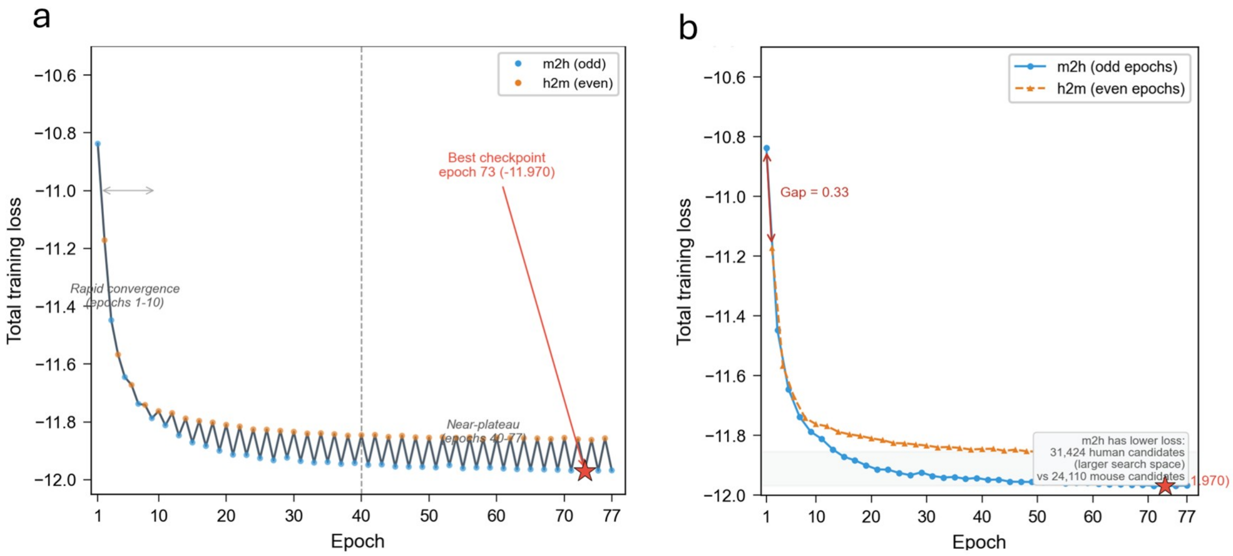


Figure 1—figure supplement 2. Training trace. (a) The displayed run covers 77 epochs and identifies epoch 73 as the best checkpoint, with total training loss of -11.970. The dashed line at epoch 40 is a reference marker. (b) Odd-epoch mouse-to-human and even-epoch human-to-mouse losses are shown separately. A single trace does not constitute a repeated-seed stability experiment or establish an identical training history for every model variant.

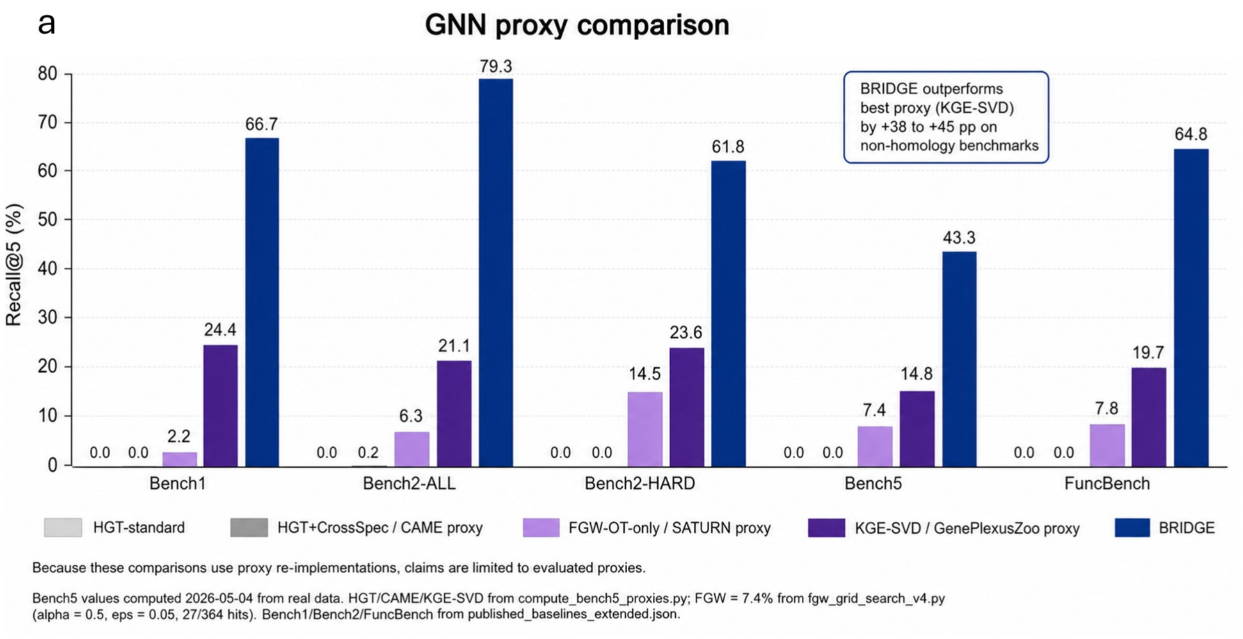


Figure 2—figure supplement 1. Standalone component comparisons. The embedding, FGW and SVD evaluations are internal study components rather than independently reproduced executions of externally named methods. Supplementary File 6 supplies the available Bench1, Bench2 and FuncBench component records; it does not contain the separate Bench5 results shown in this panel. The FGW Bench5 panel uses 364 scorable pairs out of 374, rather than an unqualified 374-pair denominator.

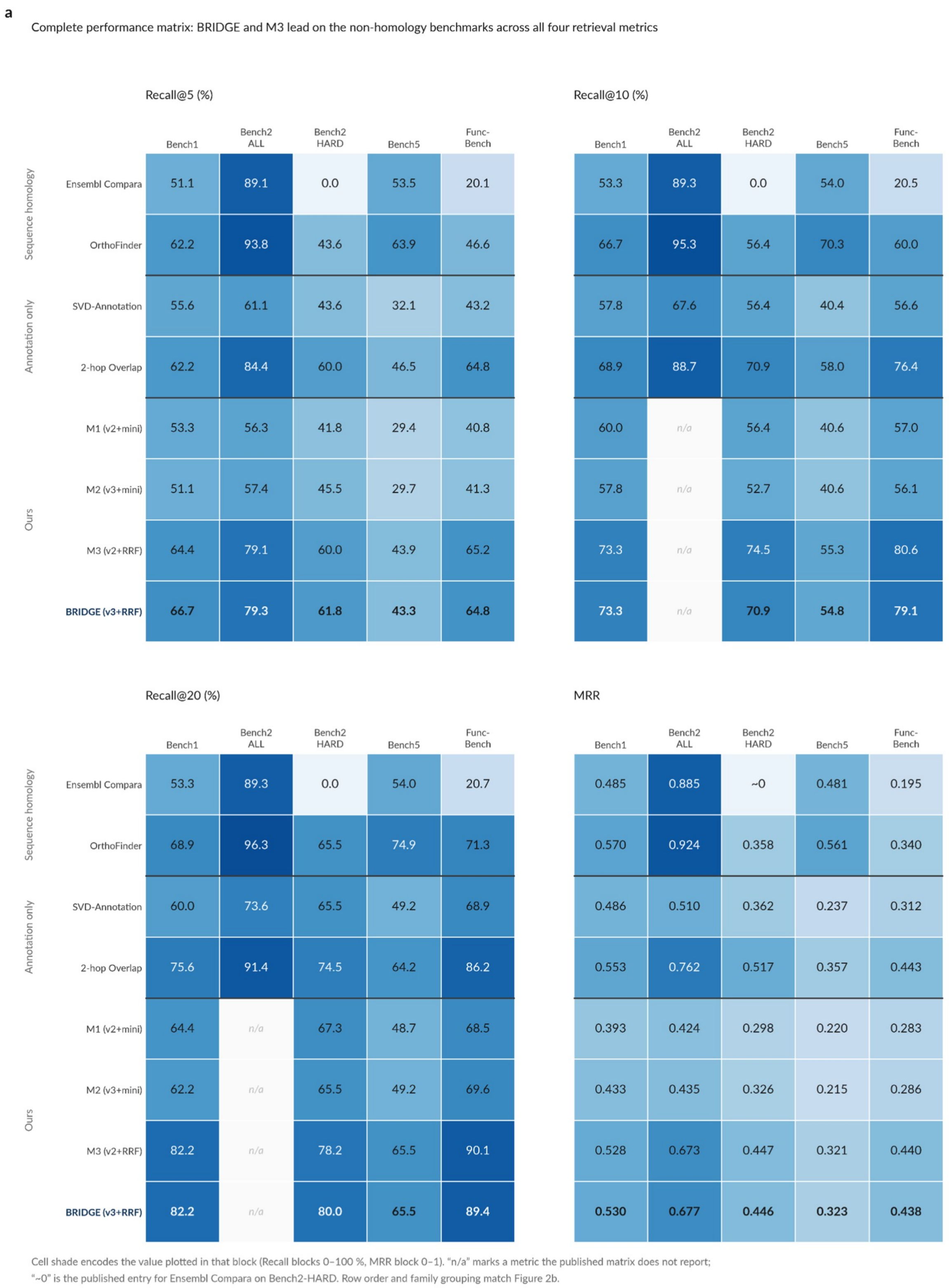


Figure 2—figure supplement 2. Complete performance matrix. Recall@5, Recall@10, Recall@20 and mean reciprocal rank are shown for the displayed methods and benchmarks. Bench2-ALL uses 512 pairs. Missing M1–M4 Recall@10 and Recall@20 values for that cohort remain missing; earlier 509-pair results are not inserted into these cells. Supplementary Files 2 and 7 provide the corresponding method and summary metrics. The OrthoFinder-labelled study comparator is described in Methods.

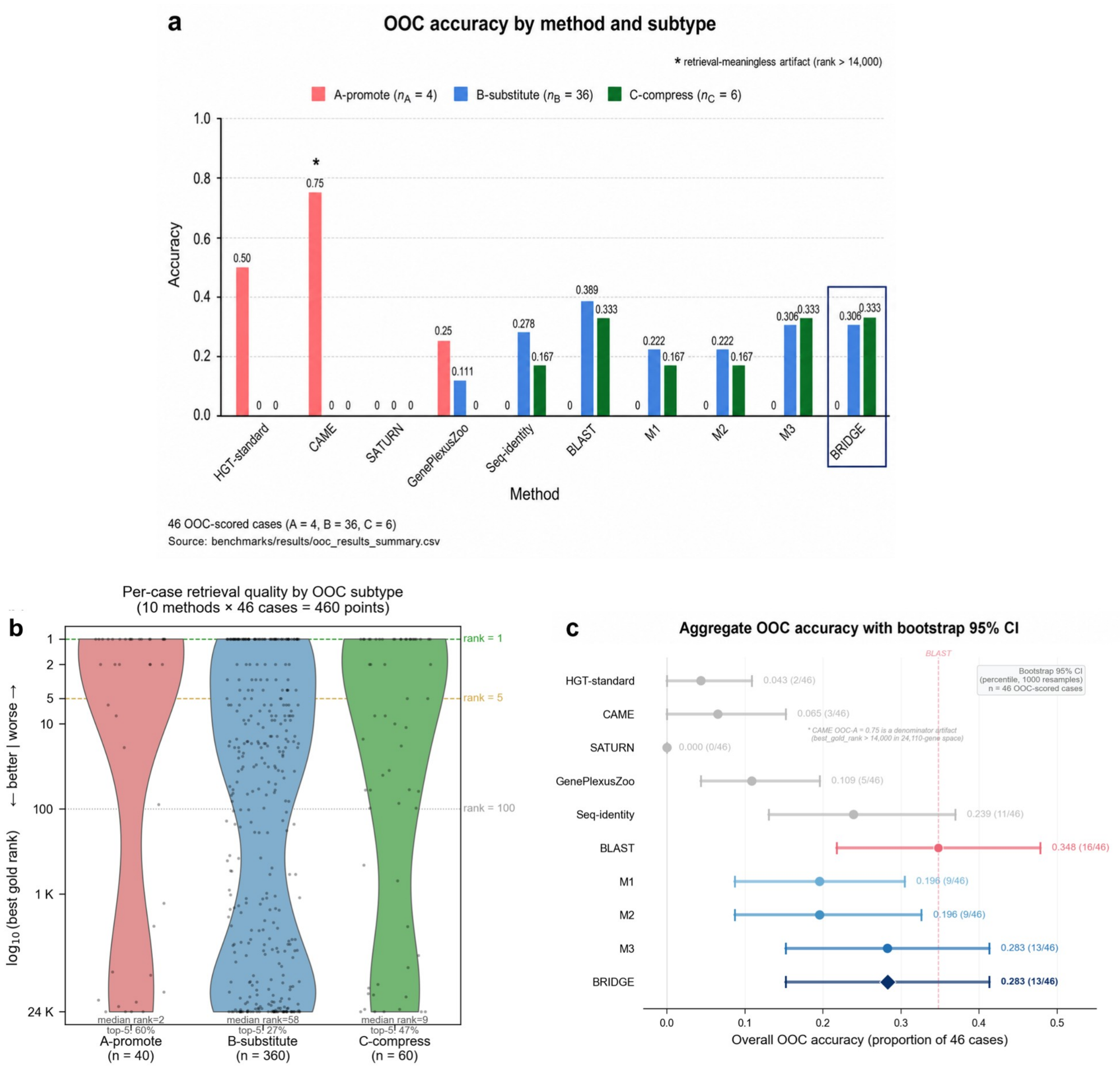


Figure 3—figure supplement 1. Ortholog-Override Criterion. The separately scored M4 evaluation contains 46 pass/fail cases: four A-promote, 36 B-substitute and six C-compress cases. Displayed passes are 0, 11 and 2, respectively. The complete roster contains 82 cases, of which 71 have a recorded M4 gold rank. Supplementary File 4 retains OOC-run values separately from the Figure 3 illustrations. The CXCL8 illustration uses a different configuration; the aggregate uses only the OOC evaluation.

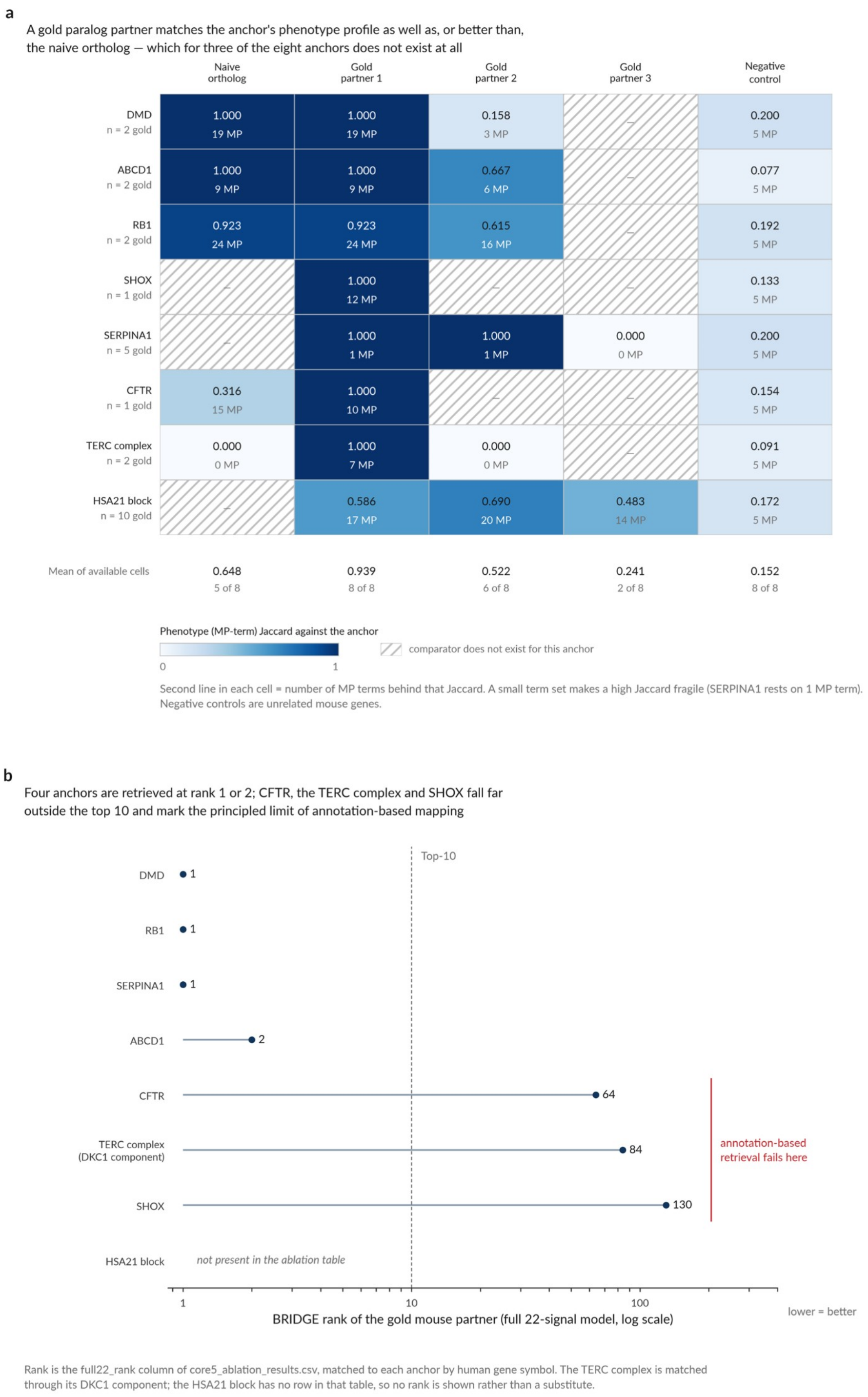


Figure 3—figure supplement 2. Phenotype concordance and legacy retrieval ranks for anchor cases. (a) MP-term Jaccard similarities compare the curated mouse partners, the naive ortholog where available and unrelated negative controls across eight anchors. Term counts are shown separately; a high similarity supported by few terms requires caution. (b) Seven anchors have matching entries in the separate 53-case ranking-strategy analysis in Supplementary File 1. The TERC complex is matched through DKC1; the HSA21 block has no matching table entry and no rank is supplied. The displayed full22 ranks are configuration-specific, not final M4 predictions. The sentinel rank 24,110 elsewhere in restricted-ranking outputs denotes an unavailable or outside-list assignment rather than a measured ordinal position in a short candidate list.

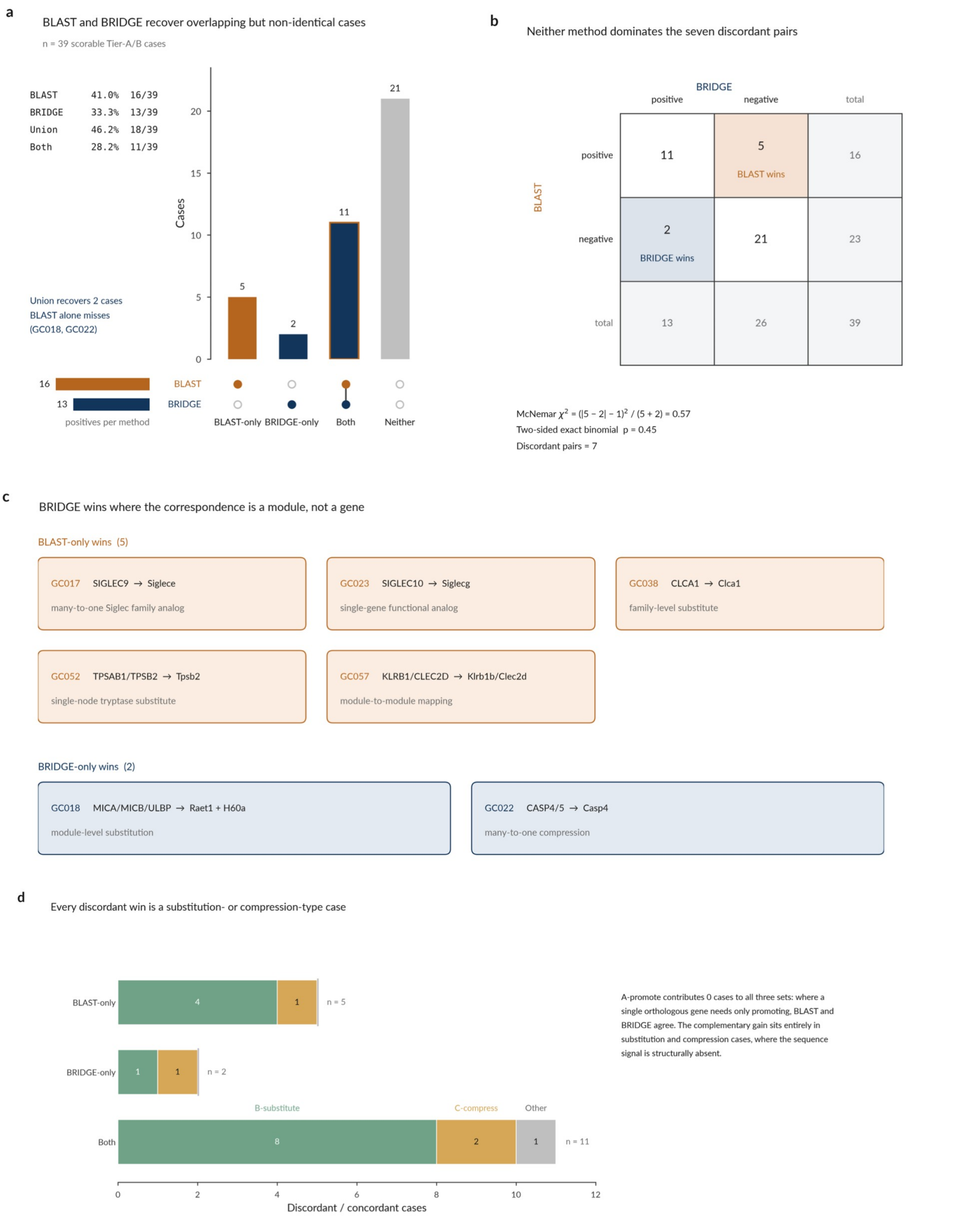


Figure 4—figure supplement 1. Complementarity on the 39-case OOC subset. The displayed baseline, M4 and union pass counts are 16, 13 and 18. Eleven cases pass both methods, five only the baseline and two only M4. The reported exact McNemar P = 0.45 does not establish a difference. The comparator labelled BLAST is the study comparator described in Methods, not a verified independent sequence-search execution.

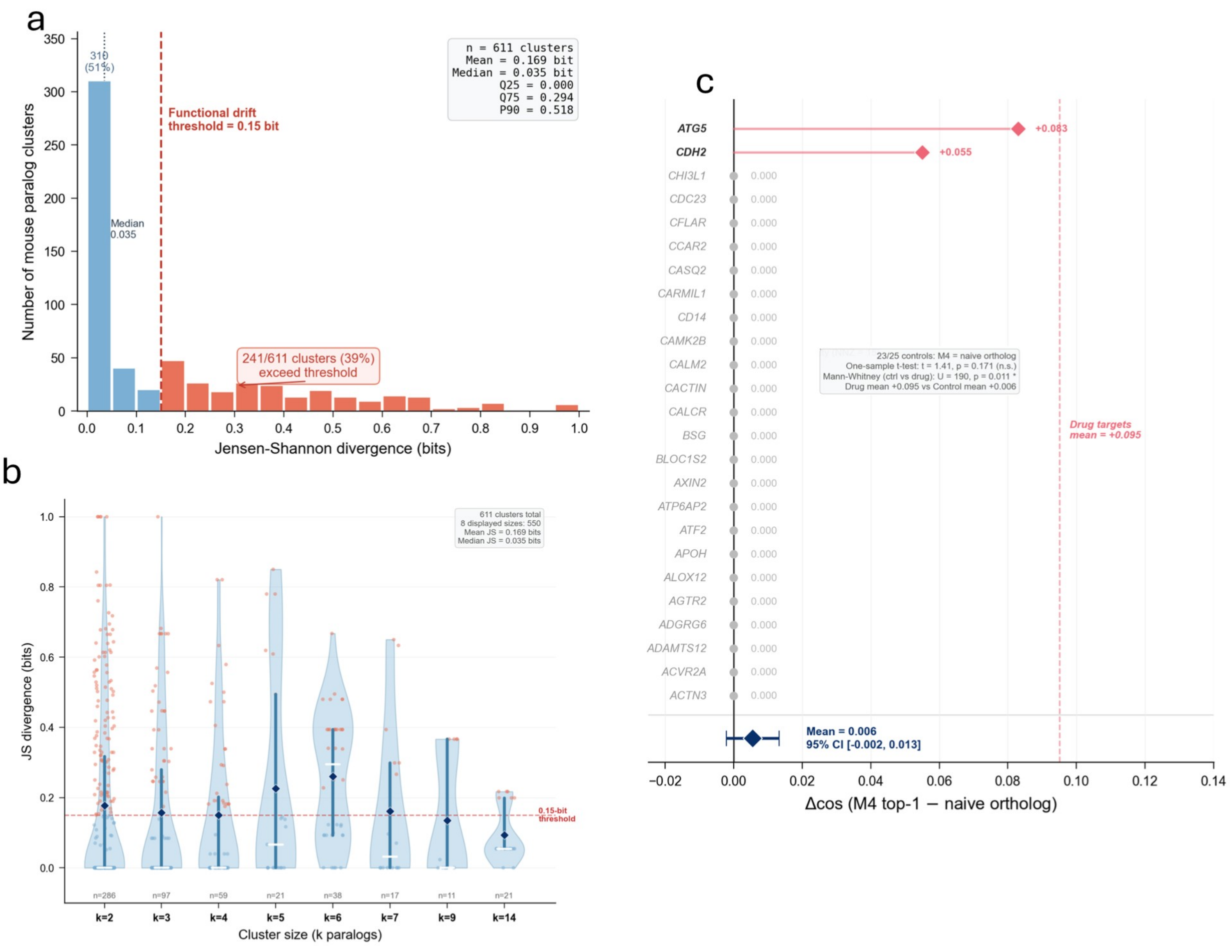


Figure 4—figure supplement 2. Annotation divergence within 611 paralog clusters. The displayed mean Jensen–Shannon divergence is 0.169 bit and the median is 0.035 bit. Of the 611 clusters, 241 exceed 0.15 bit. The panel also identifies negative controls. This describes differences in recorded annotations, which may reflect unequal evidence coverage as well as biological differences.

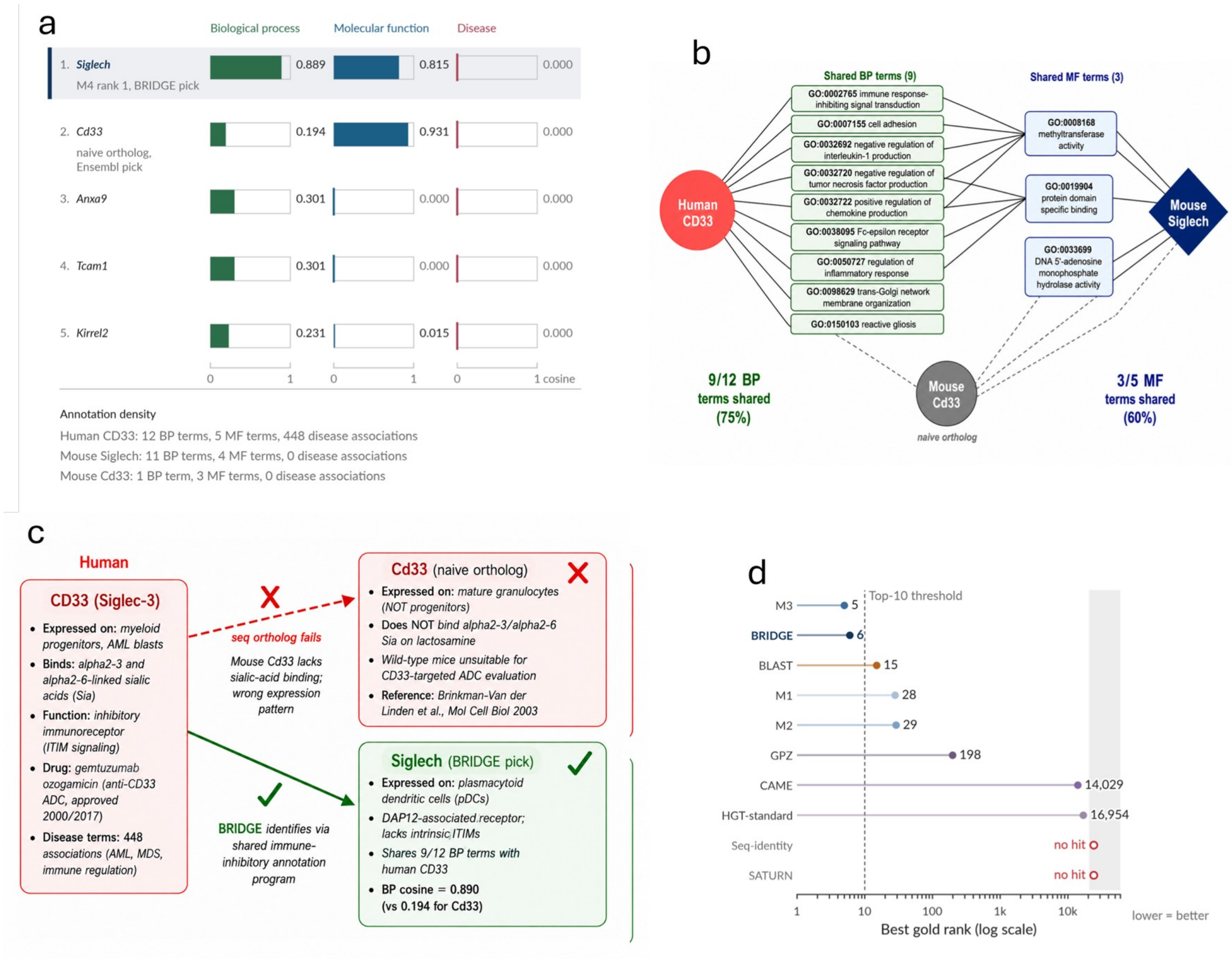


Figure 4—figure supplement 3. CD33-related candidate comparison. (a) Siglech is the BP-nearest candidate; its displayed BP and MF similarities are 0.889 and 0.815, compared with 0.194 and 0.931 for Cd33. (b) The panel displays shared GO annotations. Annotation overlap is not a biochemical or pharmacological validation. (c) The receptor-context schematic distinguishes human CD33 from mouse candidates. Siglec-H lacks intrinsic ITIMs and associates with DAP12 [18]. (d) The M4 drug-case target Cd33 has rank 6, a different candidate and ranking context from the Siglech illustration (Supplementary File 5). Candidate designation and diagram symbols do not establish CD33-like signalling or binding by an anti-CD33 therapeutic. The Siglech BP-cosine labels of 0.889 in panel a and 0.890 in panel c reflect different display rounding.

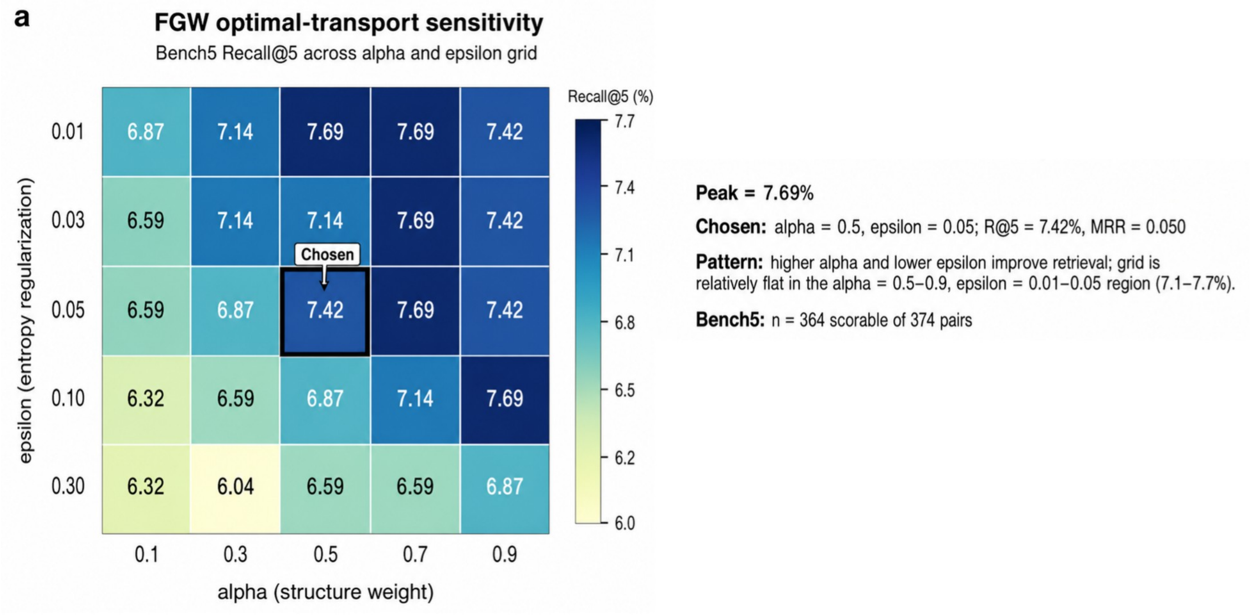


Figure 5—figure supplement 1. FGW parameter sensitivity. The grid varies the structure–feature trade-off and entropic regularization. The selected α = 0.5 and ε = 0.05 give Bench5 Recall@5 of 7.42%, compared with a grid maximum of 7.69%. This panel uses 364 scorable pairs out of the 374-pair benchmark and does not establish that the selected configuration is optimal.

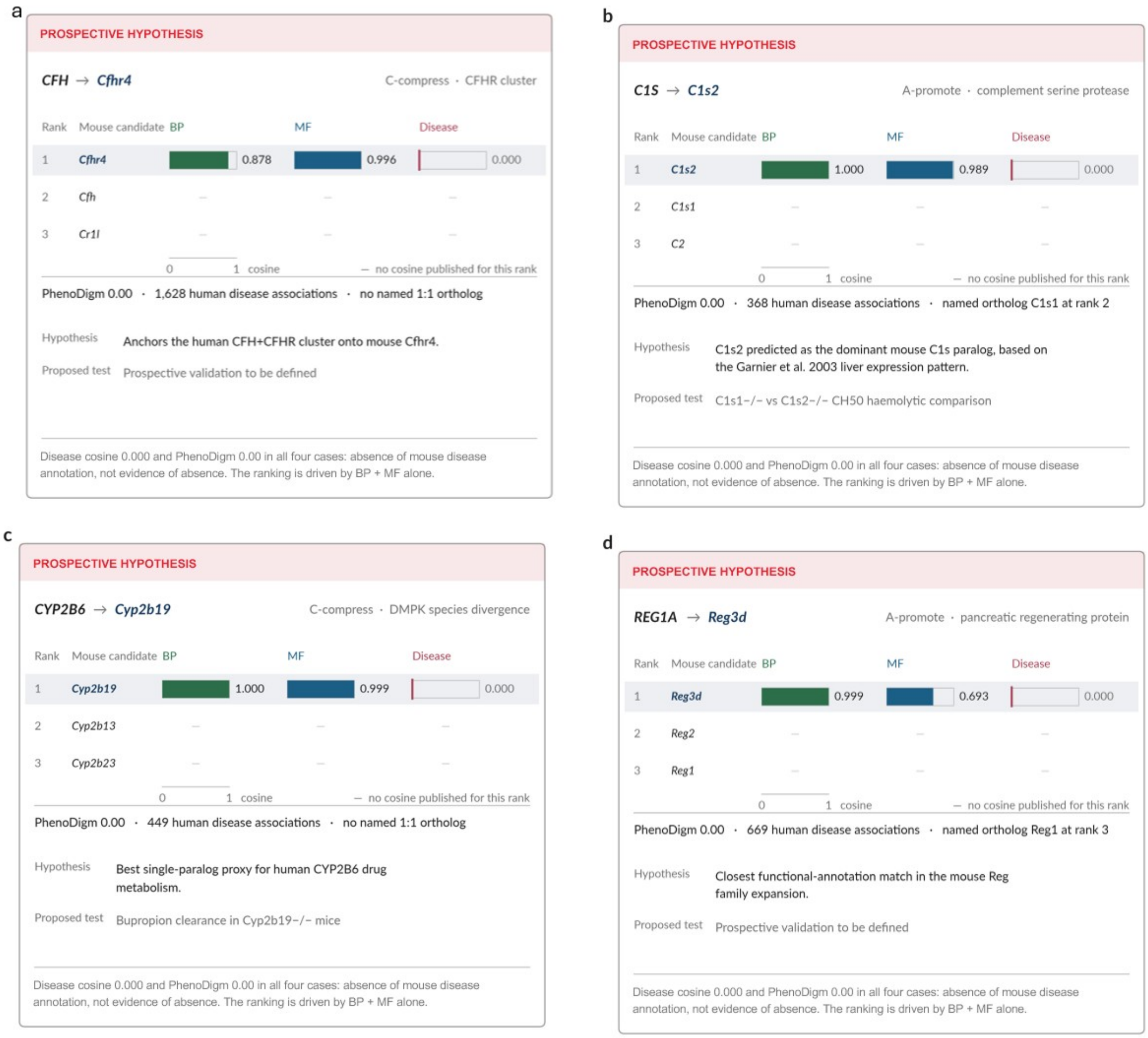


Figure 6—figure supplement 1. Four candidate functional correspondences. The panels show (a) CFH–Cfhr4, (b) C1S–C1s2, (c) CYP2B6–Cyp2b19 and (d) REG1A–Reg3d. Each panel separates BP, MF and disease annotation similarities and presents a proposed biological follow-up. The C1S shortlist record and candidate annotation evidence are provided in Supplementary Files 8 and 9; these catalogues cover a broader candidate set, not all four illustrated pairs. In Supplementary File 9, illustrated_panel links the displayed C1S records to panel b; candidate records for POT1 and APOL1 remain in the table without a current-panel reference. REG1A–Reg3d also occurs in FuncBench (Supplementary File 3), so these examples are not an entirely benchmark-independent validation set. A zero disease-similarity or phenotype score does not establish absence of disease relevance.
